# An ancient dental signalling centre supports homology of an enamel knot in sharks

**DOI:** 10.1101/2021.08.13.456270

**Authors:** Alexandre P. Thiery, Ariane S. I. Standing, Rory L. Cooper, Gareth J. Fraser

## Abstract

Development of tooth cusps is regulated by the enamel knot signalling centre. Fgf signalling regulates differential proliferation between the enamel knot and adjacent dental epithelia during tooth development, leading to formation of the dental cusp. The presence of an enamel knot in non-mammalian vertebrates is debated given differences in signalling. Here we show the conservation and restriction of *fgf10* and *fgf3* to the sites of future dental cusps in the shark (*Scyliorhinus canicula*), whilst also highlighting striking differences between the shark and mouse. We reveal shifts in tooth size, shape and cusp number following small molecule perturbations of canonical Wnt signalling. Resulting tooth phenotypes mirror observed effects in mammals, where canonical Wnt has been implicated as an upstream regulator of enamel knot signalling. *In silico* modelling of shark dental morphogenesis demonstrates how subtle changes in activatory and inhibitory signals can alter tooth shape, resembling phenotypes observed following experimental Wnt perturbation. Our results support the functional conservation of an enamel knot-like signalling centre throughout vertebrates and suggest that varied tooth types from sharks to mammals follow a similar developmental bauplan. Lineage-specific differences in signalling are not sufficient in refuting homology of this signalling centre, which is likely older than teeth themselves.

## Introduction

Diversification of the dentition has been instrumental in the success of vertebrates. It has become highly adapted to its respective environmental niche, and as a result has given rise to a plethora of unusual forms. It is generally observed that there is an overall increase in dental morphological complexity throughout evolution, culminating in mammals which possess multiple regionalised tooth types (heterodonty) [1,2]. A trade-off between tooth morphological complexity and dental regenerative ability is thought to exist within the mammalian lineage, with precise occlusion favoured at the expense of dental regeneration [1]. The study of dental development has mostly focussed on transcriptional regulators, which mediate growth, morphogenesis and cellular differentiation. Dental development is tightly regulated via interactions between the oral epithelium and underlying neural-crest derived mesenchyme [3]. Canonical Wnt, fibroblast growth factor (Fgf), hedgehog (Hh), bone morphogenetic protein (Bmp) and Notch signalling pathways have all been implicated as important regulators of tooth development and morphogenesis [4].

Teeth develop from an initial epithelial placode, which proceeds to proliferate and grow in size during the subsequent bud stage. Following the bud stage, the tooth enters the cap stage, which marks the onset of morphogenesis. Morphogenesis of epithelial appendages coincides with a change in shape of the epithelial placode. This process can be driven via differential proliferation rates (i.e., teeth [5,6], cell shape changes (i.e., intestinal crypt [7], or cell migration (i.e., hair [8]). Signalling molecules drive reciprocal epithelial to mesenchymal signalling and regulate the morphogenesis of epithelial appendages. Canonical Wnt, fibroblast growth factor (Fgf), hedgehog (Hh), bone morphogenetic protein (Bmp) and Notch signalling pathways have all been implicated in this process [4,9].

During morphogenesis the tooth acquires its defining morphological feature: the dental cusp. In mammals, dental cusps are regulated by an epithelial signalling centre found at the tip of the first cusp, known as the primary enamel knot (EK). The EK is molecularly identifiable as non-proliferative and expresses Wnt, Bmp, Fgf and Hh markers [1]. Fgf signalling drives differential proliferation between the EK and rapidly proliferating adjacent dental epithelium, leading to folding of the epithelium at the site of dental cusps and the formation of the final tooth shape [5,6,10]. During the final stages of dental morphogenesis, the EK dramatically reduces in size as cells undergo apoptosis [10]. In multi-cuspid teeth, such as mammalian molars, subsequent signalling centres termed secondary enamel knots (SEK) form at the site of each extra cusp and are thought to be induced by the EK. Folding of the dental epithelium between primary and secondary EKs leads to formation of cervical loops in-between each cusp [6,11]. Most of our understanding of tooth shape comes directly from the study of mammalian molar cusp development. Whilst it is known that EK signalling centres are found throughout mammals, their presence in other vertebrates is less clear.

Whilst over a dozen markers have been described within the mammalian EK, Wnt signalling appears to be a primary upstream regulator of EK signalling. Lef1-/- mutants exhibit arrested tooth development during bud stage, with the tooth cap and associated cusps failing to both form and express EK markers, including Fgf4, Shh and Bmp4 [12]. Fgf4 is capable of rescuing tooth development in Lef1-/- mutants [12]. Furthermore, the inhibition of Wnt signalling during early bud stage via ectopic expression of the Wnt antagonist Dickkopf-related protein 1 (Dkk1), leads to blunted cusps and a reduction in Bmp4 signalling [13]. These results suggest that Wnt signalling falls upstream of Fgf and Bmp signalling in the EK. However, more complex feedback loops are involved, as mesenchymal specific knock-down of Bmp4 (Bmp4ncko/ncko) also leads to a reduction in *Lef1* expression [14].

In the subset of reptiles studied to date, there appears to be no clear histologically definable EK [15,16], although in some species there is a thickening of the inner dental epithelium which leads to the asymmetric deposition of enamel and the formation of cusps [17]. Furthermore, whilst in reptiles, cells of the inner dental epithelium are non-proliferative, this region is not as highly restricted to the cusp as in mammalian molars [15,18]. However, there is conservation of signalling within an EK-like signalling centre [19]. Similar EK-like signalling centres have been described in teleosts, with chemical perturbation of these signalling pathways resulting in shifts in cusp number [20]. Sharks, which are basal crown gnathostomes, also possess a variety of tooth types, with clearly defined cusps. In sharks and rays, there is a region of non-proliferative cells within the apical dental epithelium thought to be associated with the primary cusp (i.e., the primary enameloid knot; [21,22], however the presence of a definitive EK has not yet reached a consensus [23], and some have even argued that the EK is uniquely a mammalian innovation [16,18,19,24].

Sharks and rays (elasmobranchs) represent two major subdivisions of the cartilaginous fishes. The diversity in tooth shape in the elasmobranchs is vast, ranging from flattened tooth units associated with crushing prey items, to multi-cuspid and serrated tooth units necessary for cutting [25]. The presence of multiple cusps as a standard tooth form for most sharks suggests that is conserved developmental control of tooth cusp morphogenesis across vertebrates. Given the conservation of general tooth development among vertebrates [16,21,22,26–28], it seems appropriate that the epithelial and mesenchymal signals regulating dental morphogenesis would also be conserved, including the EK signalling centre. This would imply that this crucial tooth signalling centre is older than the mammalian clade. Therefore, due to the vast array of dental phenotypes in sharks and their position in the wider context of vertebrate phylogeny, we employ the shark as a developmental model for non-mammalian tooth morphogenesis.

In order to determine the extent of signalling conservation between mammals and sharks during dental morphogenesis, we documented the expression of mammalian EK markers in the small-spotted catshark, *Scyliorhinus canicula*. Furthermore, given the importance of Wnt signalling during dental morphogenesis and cusp formation in mammals, we sought to perturb Wnt signalling and determine its function during catshark tooth development. Following small-molecule Wnt-signalling perturbation, we identify shifts in tooth shape, size and cusp number, and model the resulting phenotypes *in silico* using the ‘ToothMaker’ programme [5]. Our results reveal that despite differences in molecular signalling between the catshark and mammals, the fundamental components of the enamel knot signalling centre are highly conserved. Therefore, EK signalling at the apex of the developing tooth cusp is a vertebrate innovation and contributes to the vast range of vertebrate dental phenotypes.

## Results

### Histological and morphological analysis of the catshark dentition

The first dental generation develops relatively superficially on the oral surface before the dental lamina has fully invaginated (Figure 1D). As the dental lamina grows, more generations emerge. New teeth are initiated at the tip of the dental lamina and move anteriorly in a conveyor belt-like manner. As teeth move along this trajectory, they undergo morphogenesis and matrix secretion, before erupting on the oral surface on the labial side of the dental lamina (Figure 1E). Adjacent tooth families are staggered in the timing of their initiation (Figure 1C), resulting in differences in the developmental stages of adjacent teeth.

**Figure 1.**
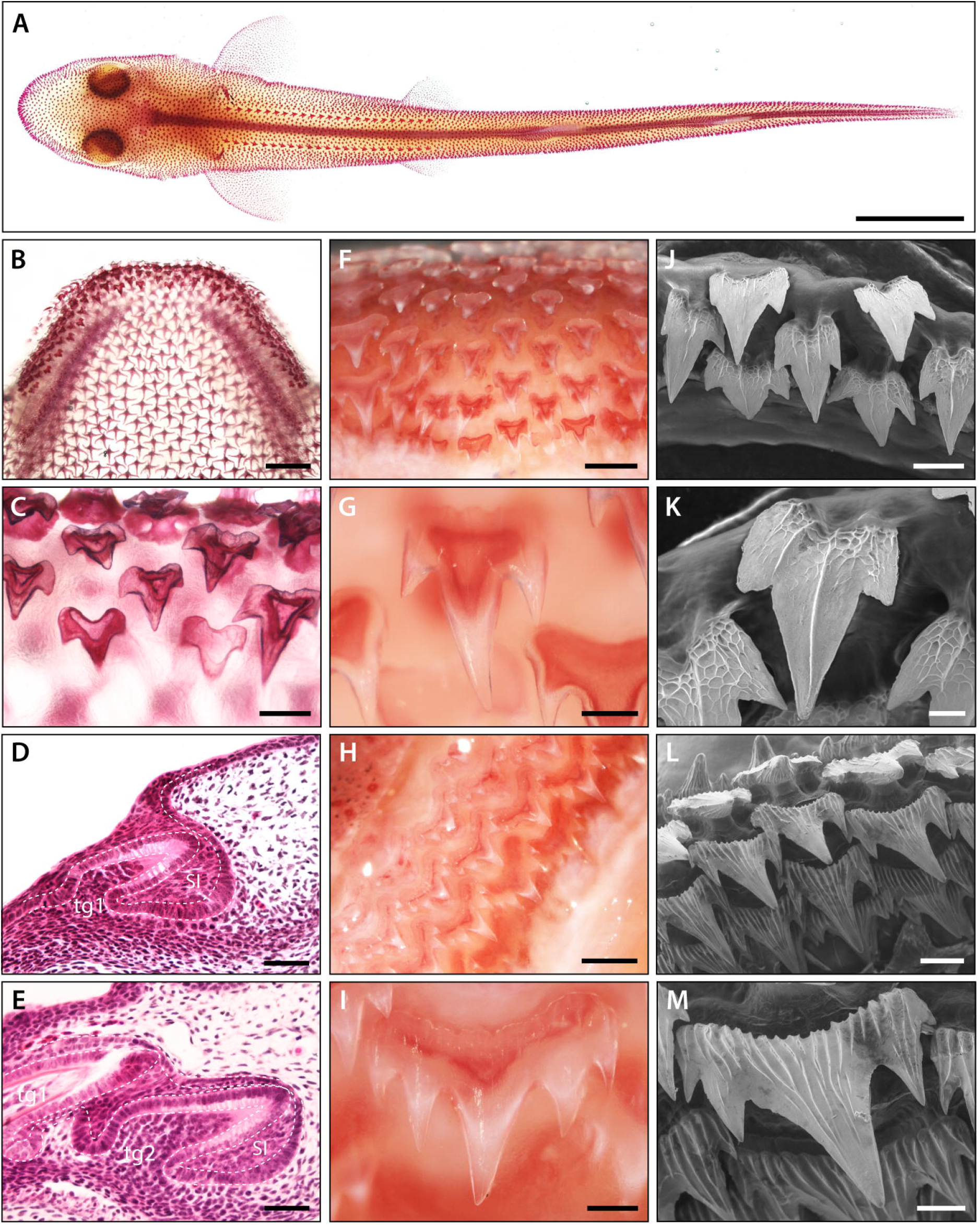
Pattern and morphology of the catshark dentition. Images of catshark samples cleared and stained with alizarin red, reveal pattern and morphology of the dentition in the lower jaw (A-C and F-I). A) Dorsal view of hatchling stage catshark (Stage 34) revealing denticle on the body surface. B) Dorsal view of hatchling stage (Stage 34) lower jaw. C) Magnification of B, showing staggered pattern of adjacent tooth families. Histological staining with haematoxylin and eosin on sagittal cross sections through early stage 32 (D) and late stage 32 (E) lower jaws shows the growth of the DL during dental development. The first dental generation (tg1) develop relatively superficially at the oral surface, before full invagination and elongation of the DL (D). The DL then grows deep into the underlying mesenchyme, with the second (tg2) and subsequent dental generations initiated at the SL (E). The addition of numerous successional dental generations can be seen in the adult jaw (F and H). At the jaw symphysis (F), teeth often remain tricuspid (G). However, in lateral regions (H), teeth develop 5-7 cusps (I). Scanning electron microscope (SEM) images reveal tricuspid teeth in the embryo (stage 33) (J-K) and pentacuspid teeth in the adult (L and M). Scale bars are 10mm in A; 1mm in B, F and H; 250μm in C, G, I and M; 50μm in D, E and K; 125μm in J and 500μm in L. ora; oral, abo; aboral, lin; lingual, lab; labial.

There is an observable shift in cusp number between embryonic and juvenile catshark stages (Figure 1J-M). Embryonic teeth generally develop with three cusps (Figure 1C, J and K). In contrast, adult catsharks can possess anywhere between three and seven cusps depending on the position of the tooth along the jaw margin (Figure 1 F-I). Given these changes in cusp number, we aimed to investigate how the development of cusps is regulated. In order to see if there is a conserved EK signalling centre in the catshark, we documented the expression of known mammalian EK markers during odontogenesis (Figure 2). In situ hybridisation was undertaken on sagittal paraffin sections to observe the expression of EK markers. We used embryonic catshark samples at late Stage 32 (~125 days post fertilisation (dpf)) [29], as two to three dental generations had already undergone the process of dental initiation by this stage, allowing us to compare early and late stage dental morphogenesis.

**Figure 2.**
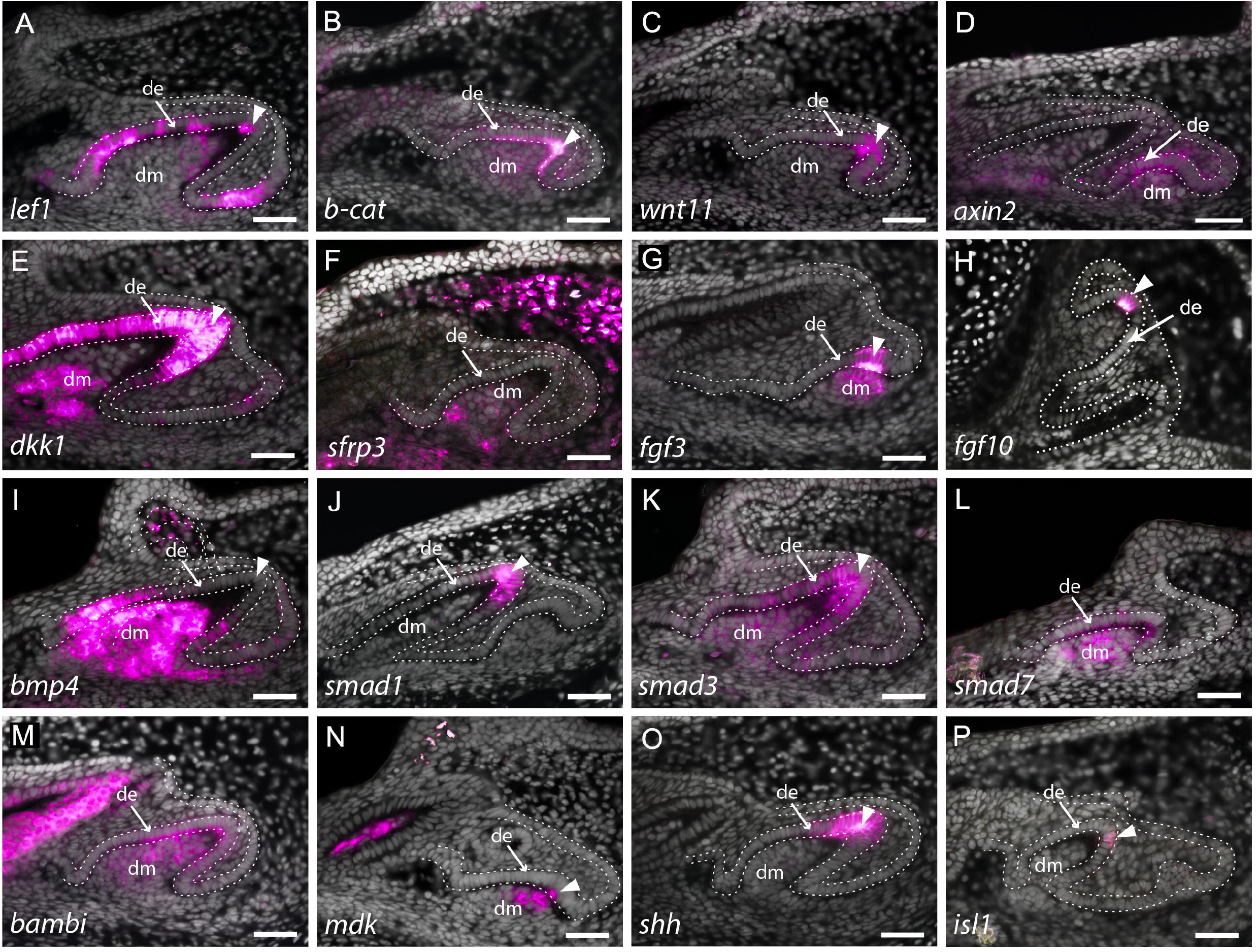
Expression of enamel knot markers during catshark dental morphogenesis. *In situ* hybridisation assay on sagittal paraffin sections of late stage 32 catshark jaws, reveal the expression of markers involved in major developmental pathways, including canonical Wnt signalling (A-F), Fgf signalling (G and H) and Bmp signalling (H-L). Wnt markers, lef1 (A), β-catenin (B) and wnt11 (C) are all expressed within the apical tip of the developing tooth during cap stage. Weak expression can also be seen within the dental mesenchyme and surrounding dental epithelium in lef1 (A) and β-catenin (B). Weak expression of the Wnt inhibitor, sfrp3 is found within the dental mesenchyme during cap stage, whereas dkk1 is also found within the dental epithelium later in morphogenesis (D). The expression of Fgf markers fgf3 (F) and fgf10 (G) is highly specific to the EK during late bud to early cap stage, with fgf3 weakly expressed in the dental mesenchyme bellow the EK. bmp4 (H) expression is absent from the EK during late morphogenesis, but is strongly expressed within both the rest of the dental epithelium and dental papilla. smad1 (I) is found within the apical tip of the dental epithelium, whilst smad3 (J) is also expressed in the dental mesenchyme during late cap stage. Bmp inhibitors smad7 (K) and bambi (L) are both expressed throughout both epithelium and mesenchyme of developing teeth. mdk (M) and isl1 (O) expression is restricted to a few cells of the EK, with mdk also expressed within the underlying dental mesenchyme. In contrast, shh (N) expression is extensive broad, but is still restricted to the apical tip of the dental epithelium. Gene expression is false coloured in magenta. White arrowhead points to expression within the apical tip of developing teeth and putative EK. White dotted lines depict the columnar basal epithelial cells of the dental lamina and dental epithelium. DAPI nuclear stain is false coloured in grey. All images are of lower jaws, except G, which is of the upper jaw. Scale bars are 50μm. de, dental epithelium; dm, dental mesenchyme.

### Key markers of tooth morphogenesis in sharks

Canonical Wnt signalling plays a crucial role in the formation of dental cusps in mammals [12,13]. In the catshark, transcription factors *Lymphoid enhancer binding factor-1* (*lef1*) (Figure 2A) and *β-catenin* (Figure 2B) are both expressed within developing bell stage teeth. Weak expression can be seen in the dental mesenchyme (Figure 2: dm) and regions of the dental epithelium (Figure 2: de). However, there is also an observable upregulation of their expression within the apical dental epithelium (Figure 2: white arrowhead). Wnt11 has been previously identified as an activator of non-canonical Wnt signalling, although its expression has also been shown to stimulate canonical Wnt signalling in a case-specific context [13]. We note the upregulation of *wnt11* specifically within the apical dental epithelium during cap stage (Figure 2C). The downstream target of canonical Wnt signalling, *axin2*, is also expressed within the early dental epithelium; however, its expression is not specifically restricted to the apex (Figure 2D).

Canonical Wnt antagonists *Dickkopf 1* (*dkk1*) and *Frizzled-related protein 3* (*sfrp3*) are also expressed within the developing tooth. *dkk1* is seen within both the dental papilla and throughout the dental epithelium in late morphogenesis (Figure 2D and 3C). Reduced proliferation is a key requirement of an EK, with differential proliferation rates within the dental epithelium leading to the formation of dental cusps [6]. Double *in situ*/immunohistochemistry for *dkk1* and proliferative cell nuclear antigen (PCNA) revealed a small number of non-proliferative cells at the tip of the tooth during cap stage (Figure 3Ba), with a marked reduction in proliferation throughout the entire dental epithelium late morphogenesis (Figure 3Ca). Although *dkk1* is expressed throughout the entire dental epithelium during late morphogenesis (Figure 2E), during cap stage its epithelial expression is specifically restricted to the non-proliferative cells corresponding to the putative EK (Figure 3B). In the pig [30], *dkk1* expression is restricted to the dental mesenchyme, with no expression observed within the dental epithelium. In contrast to *dkk1*, *sfrp3* is specifically restricted to the dental mesenchyme, with weak expression within the dental papilla (Figure 2F). In the mouse, *sfrp3* is also restricted to the mesenchyme [31], though its expression is much stronger than that observed in the shark (Figure 2F). Not all canonical Wnt-related markers are expressed in equivalent tissue types during the cap to early bell stage of dental development between mammals and the shark. However, there is expression of Wnt markers within both the dental epithelium and dental mesenchyme, with an isolated yet upregulated region of expression within the putative EK. The expression of canonical Wnt antagonists *dkk1* and *sfrp3* may also be playing a role in restricting Wnt activity within the dental mesenchyme.

**Figure 3.**
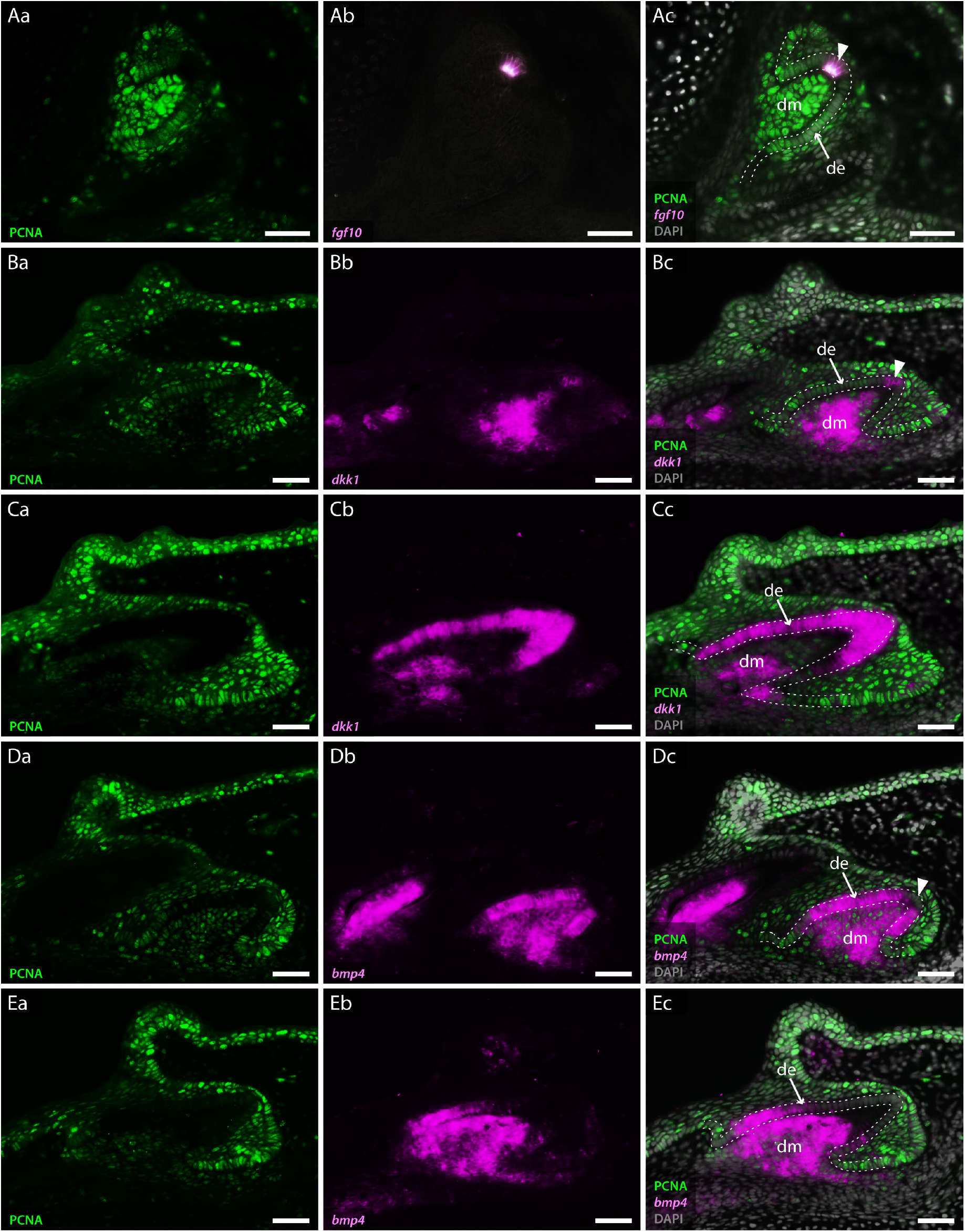
Co-expression of odontogenic markers regulating dental morphogenesis with PCNA. Double section in situ hybridisation/immunohistochemistry on catshark jaws reveals co-expression of markers involved in EK signalling and proliferative cell nuclear antigen (PCNA), within teeth undergoing cap stage (A, B and D) and late morphogenesis (C and E). Images Aa-Ea reveal expression of PCNA. Images Ab-Eb reveal expression of in situ hybridisation markers. Images Ac-Ec reveal co-expression of PCNA and in situ hybridisation markers. PCNA expression is absent from the tip of the dental epithelium, corresponding to the EK in cap stage teeth (Aa, Ba and Da). The extent of PCNA expression within developing teeth decreases as teeth undergo morphogenesis (Ca and Ea). fgf10 is expressed within a small subset of dental epithelial cells corresponding to the EK (Ab). Its expression is inversely complementary to the expression of PCNA (Ac). dkk1 expression is initially upregulated in the dental mesenchyme during cap stage (Bb), with restricted epithelial expression present only within non-proliferative cells of the EK (Bc). During late morphogenesis dkk1 expression weakens within the dental mesenchyme, with observable upregulation of its expression throughout the entire dental epithelium (Cb and Cc). Unlike dkk1 and fgf10, bmp4 is absent from the apical tip of the dental epithelium throughout dental morphogenesis (Db and Eb). fgf10 (A) is shown in the upper jaw; dkk1 (B and C) and bmp4 (D and E) are shown in the lower jaw. Gene expression is shown in magenta, PCNA protein expression is in green and DAPI nuclear stain in grey. White dotted lines depict the outer dental epithelium of the developing tooth. Scale bars are 50μm. de, dental epithelium; dm, dental mesenchyme.

Interestingly, the expression of *fgf3* and *fgf10* is almost identical between the shark and the opossum (*Monodelphis domestica*), which shows a subtle shift in expression of *Fgf10* compared to placental mammals, with *Fgf10* expressed in both the EK and the underlying dental mesenchyme in the opossum, and not in the EK of the mouse [32,33]. We find *fgf3* expressed in the tip of the dental epithelium, whilst its mesenchymal expression is confined just below the epithelium (Figure 2G). The expression of *fgf10* is highly restricted to only a few epithelial cells at the tip of the developing tooth during cap stage (Figure 2H). This expression is located precisely within a small cluster of non-proliferative dental epithelial cells (Figure 3Ab), marked by a lack of PCNA expression (Figure 3Aa). The importance of Fgf signals in regulating the differential proliferation of the EK relative to surrounding dental epithelium [6,32], together with the highly specific expression of *fgf10* within this non-proliferative region provides strong support of an EK in the catshark. The similarity in expression between marsupial mammals and sharks raises the possibility that loss of Fgf10 in the EK in placental mammals is a lineage-specific modification in EK signalling.

During the induction of mammalian molars, Bmp4 is involved in reciprocal epithelial to mesenchymal signalling, during which the odontogenic potential shifts from the dental epithelium to the dental mesenchyme. Concordant with this shift in odontogenic potential, *bmp4* shifts from epithelial and mesenchymal expression, to mesenchymal only [34]. As with *fgf3* (Figure 2G), *lef1* and *β-catenin* (Figure 2A and B) [21], *bmp4* is also expressed in the dental epithelium and condensing mesenchyme during bud stage in the catshark (Figure S1A). The increase in mesenchymal *bmp4* expression between bud (Figure S1A) and cap stages (Figure S1B), reflects the epithelial to mesenchymal shift in odontogenic potential observed in mammals [34]. However, unlike in mammals, *bmp4* remains expressed within sub-regions of the dental epithelium throughout the duration of morphogenesis (Figure 2I; S1). Given the epithelial expression of *bmp4* during bud stage, it had previously been suggested that it may play a role in putative EK signalling in the catshark [21]. Following the epithelial to mesenchymal shift in Bmp4 in the mouse, it is then secondarily upregulated within the EK during late cap stage [10]. However, we do not observe any secondary upregulation within the putative EK in the catshark. Instead, its expression is rapidly downregulated within the apical tip of the tooth (Figure 2I; Figure 3D). This region of bmp4 downregulation corresponds specifically to a restricted group of non-proliferative epithelial cells (Figure 3Dc). Throughout later stages of morphogenesis, *bmp4* becomes further restricted to only the lateral epithelial cells of the developing tooth (Figure 3E). The precise exclusion of *bmp4* from the non-proliferative tooth apex (Figure 3D and E) raises the possibility that instead of regulating putative EK signalling, it is involved in the restriction of other EK markers.

The Smad protein family plays a crucial role in TGFβ) signal transduction, including Bmp signalling [35]. Here, we show the expression of *smad1*, *smad3* and *smad7* during dental morphogenesis (Figure 2J, K, L). Smad1 and Smad3 are phosphorylated following Bmp signalling and enable its signal transduction, whilst Smad7 is a negative feedback regulator of TGFβ signalling [35,36]. *smad1* (Figure 2J) is expressed within the putative EK at the tip of the developing tooth, whilst *smad3* (Figure 2K) and *smad7* (Figure 2L) are both expressed throughout the dental epithelium and dental mesenchyme. Furthermore, as with *smad7*, BMP and activin membrane-bound inhibitor (*bambi*) is expressed in both dental epithelium and dental papilla (Figure 2M). Although interactions between these signalling molecules are complex, the expression of Bmp markers within and around the putative EK indicates a role for Bmp in EK signalling.

*fgf3*, *fgf10*, *Sonic Hedgehog* (*shh*) and *midkine* (*mdk*) have previously been observed within the apical tip of developing teeth in the catshark and little skate, corresponding to the putative EK [21,22]. Here, we also find these markers expressed within the same tissues. *mdk* is expressed within both the tooth apex and underlying dental mesenchyme (Figure 2N) whilst *shh* is found exclusively within the apical tip of the dental epithelium (Figure 2O). Another marker, *isl1*, is expressed (Figure 2P) in a very similar and restricted pattern to *fgf10*. *isl1* expression is only observed in a few apical cells associated with the putative primary EK. EKs comprise a small number of cells at the very apical tip of a developing tooth and can be hard to morphologically distinguish from surrounding dental epithelium in sagittal cross-sections. Therefore, we also used whole mount *in situ* hybridisation to examine the expression of markers throughout the entire tooth unit.

### Whole mount gene expression patterns in first generation shark teeth

As the first dental generation develops superficially, prior to full invagination of the dental lamina, teeth can be seen developing directly on the oral surface. We carried out whole mount in situ hybridisation experiments for *fgf3*, *fgf10*, *bmp4* and *shh* on early Stage 32 (~90dpf) catshark embryos, during which the first dental generation undergoes morphogenesis (Figure 4). This allowed us to identify whether markers expressed within the apical tip of developing teeth in sagittal sections were specifically restricted within cusp forming regions, in whole mount.

**Figure 4.**
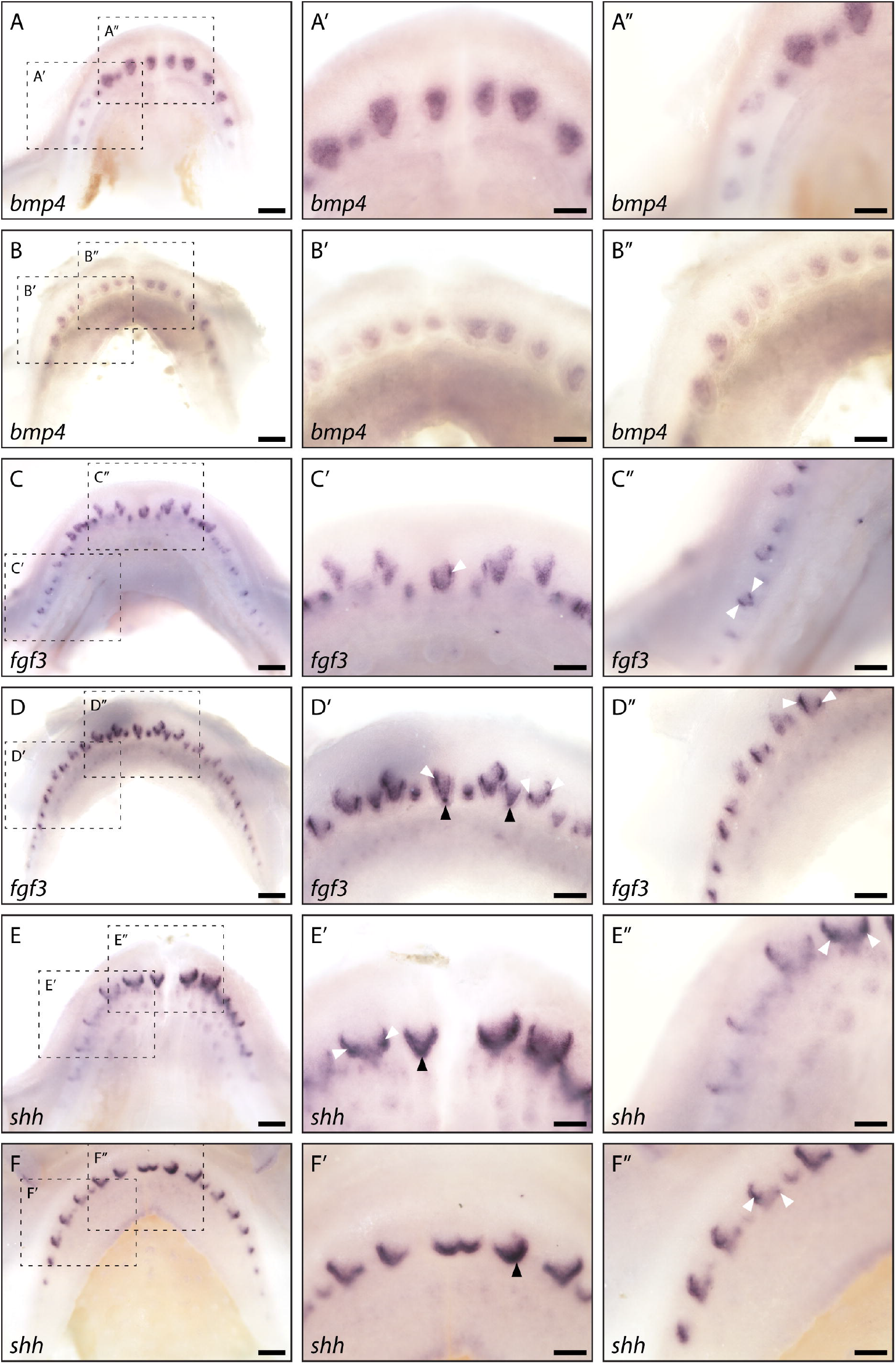
Whole mount *in situ* hybridisation reveals enamel knot-specific gene expression. Whole mount in situ hybridisation on catshark lower (A, C and E) and upper (B, D, F and G) jaws highlights the expression of bmp4 (A and B), fgf3 (C and D), shh (E and F) and fgf10 (G) across the jaw. Images are of early stage 32 (~90dpf) samples, with developing first generation teeth visible on the jaw margin. Images A-G are low magnification images. Images A’-G’ are magnified images of the central teeth along the jaw. Images A”-G” are magnified images of the left lateral side of the jaw. bmp4 expression is visible throughout the dental papilla, but appears absent from the dental epithelium (A’, A”, B’ and B”). fgf3 expression is strongly upregulated within both primary EKs (D’: black arrowheads) and secondary EKs (C’, C”, D’ and D”: white arrowheads). Weaker fgf3 expression is also noted within the dental mesenchyme (C’, C”, D’ and D”). shh expression is absent from the dental mesenchyme. However, its expression can be seen throughout the dental epithelium at the leading edge of the developing tooth (E’, E”, F’ and F”). Although its expression is not restricted to the EK throughout morphogenesis, clear shh expression within both the primary EKs (E’ and F’: black arrowheads) and secondary EKs (E’, E” and F”: white arrowheads). As with fgf3, fgf10 expression is also restricted to the future cusp forming primary (G’ and G”: black arrowheads) and secondary EKs (G’ and G”: white arrowhead), however its expression does not extend into the dental mesenchyme. Dotted black boxes in A – G depict a magnified region in A’ – G’ and A” – G”. Scale bars are 250μm in A-G and 125μm in A’ – G’ and A” – G”.

As with our section *in situ* hybridisation data (Figure 3), *bmp4* is excluded from the epithelium at the leading edge of the tooth, in whole mount. Instead, its expression appears to be primarily restricted to the dental papilla in both upper (Figure 4A) and lower jaws (Figure 4B). In contrast, *shh* can be seen expressed within the dental epithelium and specifically upregulated within the leading edge of the tooth (Figure 4E and F). *shh* is confined to the EK in mammalian molars [37,38], and although we do see strong expression within both the primary cusp (Figure 4E and F: black arrowhead) and secondary cusps (Figure 4E and F: white arrowhead), its expression is not strictly restricted at these sites. Depending on the stage of morphogenesis, the extent of *shh* expression within the inter-cusp dental epithelium is variable. Alongside playing a role in EK signalling, the expression of *shh* within the epithelium at the leading edge of the tooth suggests that it may also be involved in establishing an anterior to posterior growth gradient.

Unlike with *shh*, *fgf3* and *fgf10* expression is clearly associated with both the primary cusp (Figure 4C, D and G: black arrowhead) and secondary cusps (Figure 4C, D and G: white arrowhead). Whilst there is expression of *fgf3* within the dental mesenchyme, its epithelial expression is restricted solely to the site of future cusps. *fgf10* is exclusively expressed within the cusp forming dental epithelium, in agreement with the expression pattern observed in section (Figure 3Ab). The expression patterns of *shh*, *bmp4* and *dkk1*, in or around the dental cusps, and more importantly the precise restriction of *fgf3* and *fgf10* to these sites (Figure 4), demonstrate the conservation of an EK signalling centre at the dental cusp sites in the shark.

### Wnt-signal perturbation and geometric morphometric analysis of shark tooth morphogenesis

Wnt/β-catenin signalling has been implicated upstream of key pathways (Bmp, Fgf and Msx) regulating dental morphogenesis [12,13]. Having identified the presence of a putative EK during dental development in the catshark, we wanted to test the role of canonical Wnt signalling in the regulation of dental shape. IWR-1-endo is a canonical Wnt antagonist. It functions through stabilising cytoplasmic Axin2, in turn increasing proteasome-mediated degradation of cytoplasmic β-catenin, leading to a decrease in canonical Wnt signalling [39]. In contrast, CHIR99021 selectively inhibits the kinase activity of GSK3 [40]. This prevents the formation of the β-catenin destruction complex and in turn leads to a stabilisation of cytoplasmic β-catenin and a subsequent increase in canonical Wnt signalling [41]. Wnt signalling was both upregulated and downregulated using 2μM CHIR99021 and 1μM IWR-1-endo, respectively. We treated samples for 14 days, allowing for the initiation of, on average, one tooth generation. Samples were treated at approximately 100dpf (mid-stage 32), by which point one to two generations are undergoing morphogenesis.

Following treatment, samples were left to recover for a further 28-days before we examined the treatment effect upon final tooth shape. Drastic shifts in the shape of teeth were observed following treatment (Figure 7). Abnormalities in cusp development were seen across both IWR-1-endo and CHIR99021 treated samples, however, given the presence of reduced cusps and uneven tooth edges (serrated in appearance), quantifying the number of cusps was not feasible. In order to accurately compare changes in tooth shape following treatment, we carried out two-dimensional (2-D) geometric morphometric measurements of mineralised teeth following treatment (Figure 6).

**Figure 5.**
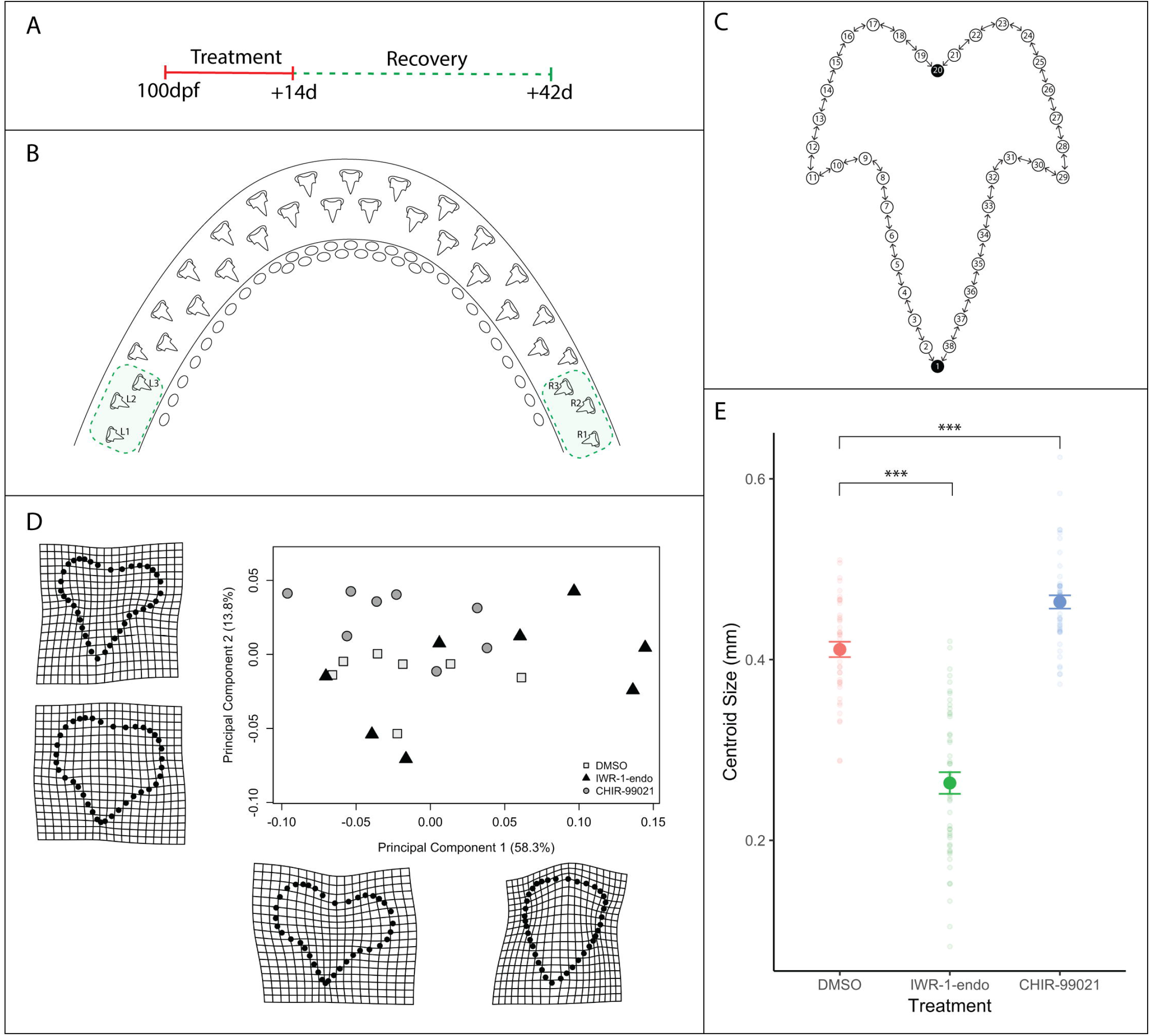
Geometric morphometric analysis reveals change in shape and size following canonical Wnt manipulation. Small molecule treatments consisted of a two-week treatment with 0.1% DMSO (control), 1μM IWR-1-endo and 2μM CHIR99021 and a subsequent four-week recovery (A). The three most lateral teeth at both left and right lower jaw margins (B) were included in the geometric morphometric analysis. A total of 38 landmarks were used to label tooth shape (C). Two fixed landmarks were placed at the tip of the primary cusp and at the base of the tooth, represented by black circles. 18 sliding semi-landmarks were placed on either side of the tooth (white circles), which were allowed to move relative to adjacent landmarks. The direction of movement is depicted by directional arrows in the schematic (C). Following Procrustes alignment of landmark coordinates and principal component analysis of the resulting shapes, the average PC1 and PC2 scores for each sample were plotted in order to depict the position of treated samples within a given shape space (D). PC1 accounted for 58.3% of the variation observed between samples, whereas PC2 accounted for 13.8%. Warpgrids shown in D reveal representative shapes at maximum and minimum PC1 and PC2 values. There is a significant effect of treatment on overall tooth shape (Procrustes ANOVA: R2 = 0.09174, F2,115 = 8.2771, p < 0.001), whilst controlling for variation within samples. There is also a significant effect of sample on shape (Procrustes ANOVA: R2 = 0.27092, F20,115 = 2.4442, p <0.001). Aside from shape, Centroid size was also measured following treatment (E). There is a significant effect of treatment on Centroid size (ANOVA: F2,115 = 235.5886, p <0.001), whilst controlling for variation within samples. There is also a significant effect of sample on Centroid size (ANOVA: F20,115 = 7.2957, p <0.001). Faint circular points are plots of each individual data point, revealing the distribution of the data. Error bars represent standard error.

**Figure 6.**
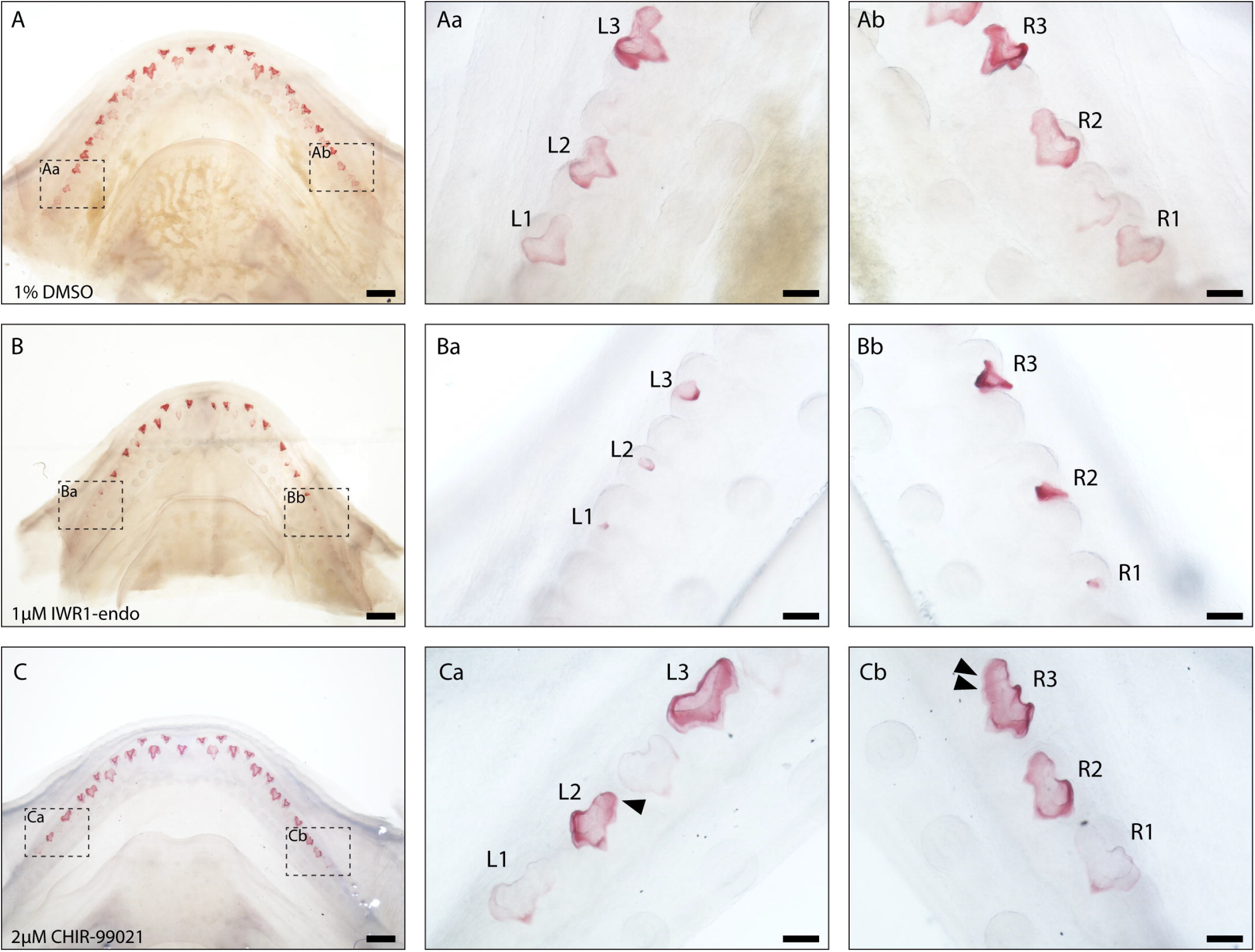
Clear and stained samples reveal observable shift in tooth shape in lateral teeth following canonical Wnt manipulation. Representative images of 0.1% DMSO (control) (A), 1μM IWR-1-endo (B) and 2μM CHIR9902 (C) treated lower jaws following two-week treatment and four-week recovery. Samples have been clear and stained. Aa-Ca and Ab-Cb are magnified images of left and right lateral jaw regions respectively. A tricuspid dentition is visible in 0.1% DMSO treated teeth (Aa and Ab). A shift to a unicuspid morphology takes place following treatment with 1μM IWR-1-endo (Ba and Bb). In contrast, a widening of the teeth is observed following 2μM CHIR9902 (Ca and Cc). Black arrowheads represent the addition of putative supernumerary cusps. Dotted black boxes in A-C depict a magnified region in Aa – Ca and Ab – Cb. Scale bars are 500μm in A-C and 100μm in Aa – Ca and Ab – Cb.

**Figure 7.**
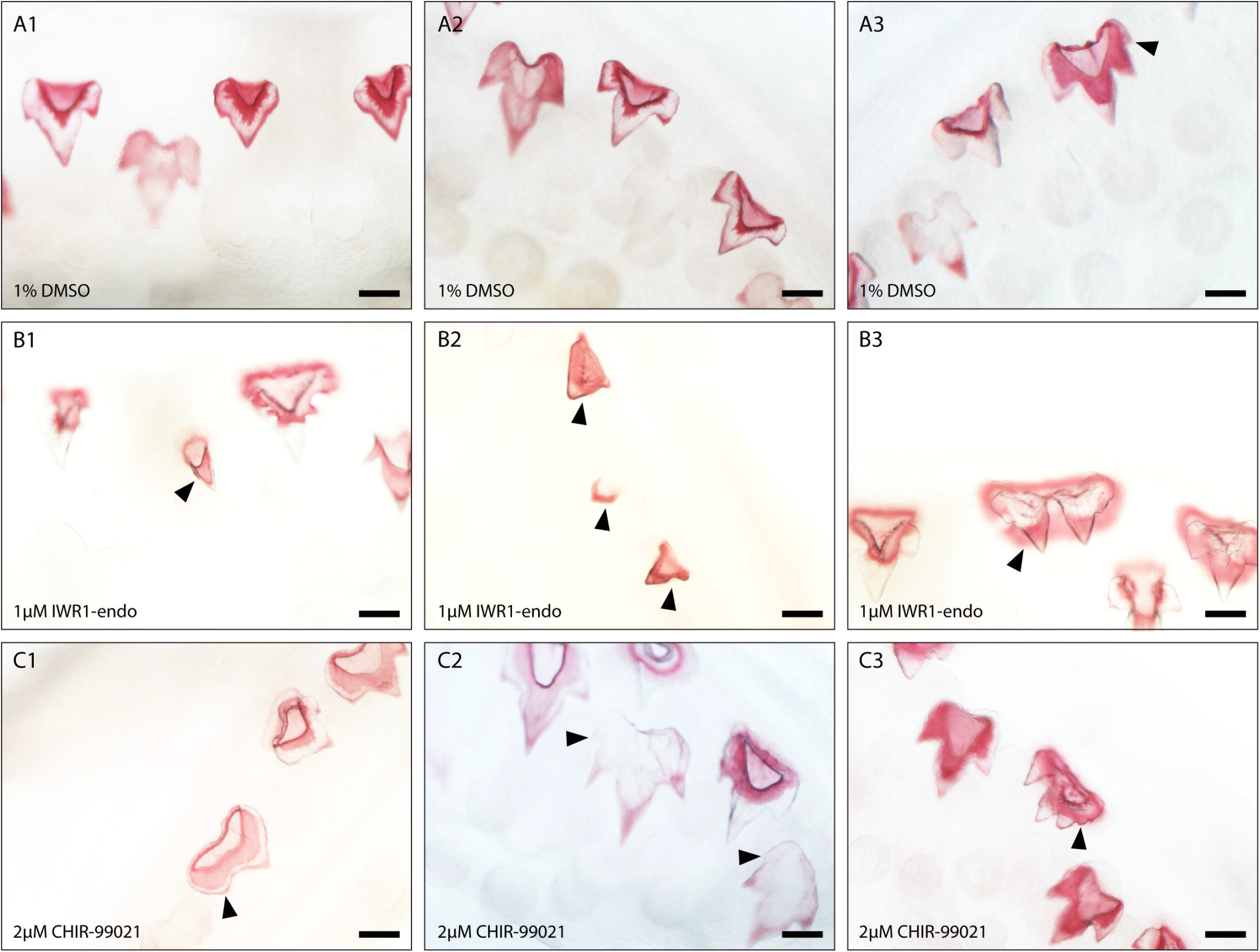
Dental diversity following canonical Wnt manipulation. Selected images depicting dental diversity following 0.1% DMSO (control) (A1-A3), 1μM IWR-1-endo (B1 B3) and 2μM CHIR9902 (C1-C3) treated lower jaws following two-week treatment and four-week recovery. 0.1% DMSO are relatively similar in shape, although the presence of four cusps is observed in a small subset of teeth (A3). Following 1μM IWR-1-endo treatment, there are numerous mineralised unicuspid and stunted teeth (B1 and B2). There are also teeth which appear duplicated in nature, but which are connected to a single root (B3). In contrast, 2μM CHIR9902 treatment leads to a widening of the teeth and defects in the position of cups (C1) and the development of supernumerary cusps (C2 and C3). Black arrowheads point to cusp defects and/or shifts from the typical tricuspid dental morphology. Scale bars are 100μm.

Landmark-based geometric morphometrics uses Cartesian coordinates instead of traditional linear measurements in order to accurately measure biological shape [42]. A measure of size known as the Centroid size can then be obtained from these coordinates. This is calculated as the square root of the sum of squared distances from each landmark to the centroid [43]. The coordinates obtained from the landmarks are then transposed, scaled (relative to the centroid size) and rotated, giving a final measurement of shape which can be compared across different treatments [43]. For our analysis, we used a 2-D landmark based geometric morphometric approach. We placed two fixed homologous landmarks on each tooth, one at the tip of the primary cusp and one at the base of the tooth. 18 sliding-semi-landmarks were then distributed evenly between these two points on either side of the tooth, given that there were no other homologous features shared by all samples (Figure 6C). Sliding semilandmarks are allowed to move between one another so that they best match corresponding points between other samples. The underlying assumption is that the curves along which the landmarks slide are homologous, even if the landmarks themselves are not [44].

We chose to compare the shape of the three most lateral teeth of both the left and right lower jaw margins (Figure 6B) to mitigate variation observed between dental generations elsewhere in the jaw. Furthermore, there is substantial variation in tooth number during this stage of development and therefore a direct comparison of tooth positions elsewhere in the jaw is problematic. Dental differentiation and mineralisation also take place unidirectionally, from the apex towards the base of the tooth. This wave of mineralisation could lead to further variability in measurements of final tooth shape. After carrying out a general Procrustes analysis (GPA), we found a significant effect of treatment on overall tooth shape (Procrustes ANOVA: R2 = 0.09174, F2,115 = 8.2771, p < 0.001), whilst controlling for variation within samples. We also found a significant effect of sample on shape (Procrustes ANOVA: R2 = 0.27092, F20,115 = 2.4442, p < 0.001), highlighting variation found within sample measurements.

Principal component analysis (PCA) of the GPA revealed that 58.3% of the variation in tooth shape observed between samples can be explained by a single axis of variation (PC1), with a further 13.8% explained by a second axis of variation (PC2) (Figure 6D). As a result of high variation in dental shape, secondary cusps are not fully represented in the morphometric analysis. However, their presence can somewhat be revealed through a bulging of the tooth at sites adjacent to the primary cusp (Fig 6D: warpgrids). The shift in dental shape observed across PC1 attributes to a change in width of the tooth and an apparent change in cusp number. 4 out of 8, IWR-1-endo treated samples can be seen exhibiting extreme unicuspid dental morphologies (PC1 = 0.05 ~ 0.15), whilst 5 out of 8 CHIR99021 treated samples exhibit wide teeth with more distinctive cusps (PC1 = −0.1 ~ 0). We also observe an increase in the variation of dental shapes across the PC1 axis following Wnt downregulation with IWR-1-endo (pairwise F-test: F41,47 = 0.277, p <0.001) (Figure S3). These results show that dental shape is being dramatically affected as a result of canonical Wnt manipulation, although it is difficult to discern directional changes in morphology. We observe not only shifts in shape as a result of treatment, but also dramatic changes in the size of teeth. Centroid size derived from GPA is a measure of an object’s scale. This provides more detail than directional measurements such as length or area which are affected by an object’s shape [42; 43]. Following treatment, we found a significant effect of treatment on Centroid size (ANOVA: F2,115 = 235.5886, p <0.001), whilst controlling for variation within samples (Figure 6E). However, there was also a significant effect of sample on Centroid size (ANOVA: F20,115 = 7.2957, p <0.001).

In order to compare the mean Centroid size between treatments, we carried out post hoc Tukey multiple comparisons of means test. There is a significant decrease in Centroid size (−35.9%) following IWR-1-endo treatment (Tukey: p <0.001), whilst conversely there is a significant increase in Centroid size (+12.8%) as a result of CHIR99021 treatment relative to control samples (Tukey: p < 0.001) (Figure 6E). These findings reveal a directional change in tooth shape as a result of canonical Wnt signalling manipulation, given that IWR-1-endo and CHIR99021 downregulate and upregulate Wnt signalling, respectively.

### Canonical Wnt signalling during development of dental cusps

Representative images of lateral teeth included in the geometric morphometric analysis reveal clear changes in tooth size (Figure 7). IWR-1-endo treatment resulted in stunted teeth, with the developed teeth failing to undergo normal morphogenesis yet successfully undergoing mineralisation (Figure 7B). Most lateral teeth exhibit a reduction in secondary cusp size, with some teeth exhibiting a unicuspid morphology with a complete loss of secondary cusps (Figure 7B). Observations of dental morphology at other sites along the jaw reveal a range of dental defects following treatment (Figure 8). There is a consistent loss/reduction in secondary cusps along the jaw (Figure 8B1 and 8B2), which is concurrent with the morphologies observed in the lateral teeth included in the geometric morphometric analysis, although not all teeth exhibit this phenotype (Figure 8B3). We also observed the development of duplicated teeth (Figure 8B3). This may arise from fusion of tooth buds belonging to adjacent tooth families, as a result of a loss in the zone of inhibition between tooth sites. However, given that adjacent teeth are staggered in the timing of their development, it is more likely that shifts in signalling during early morphogenesis has resulted in defects in the folding of the dental epithelium resulting in the formation of two primary cusps.

**Figure 8.**
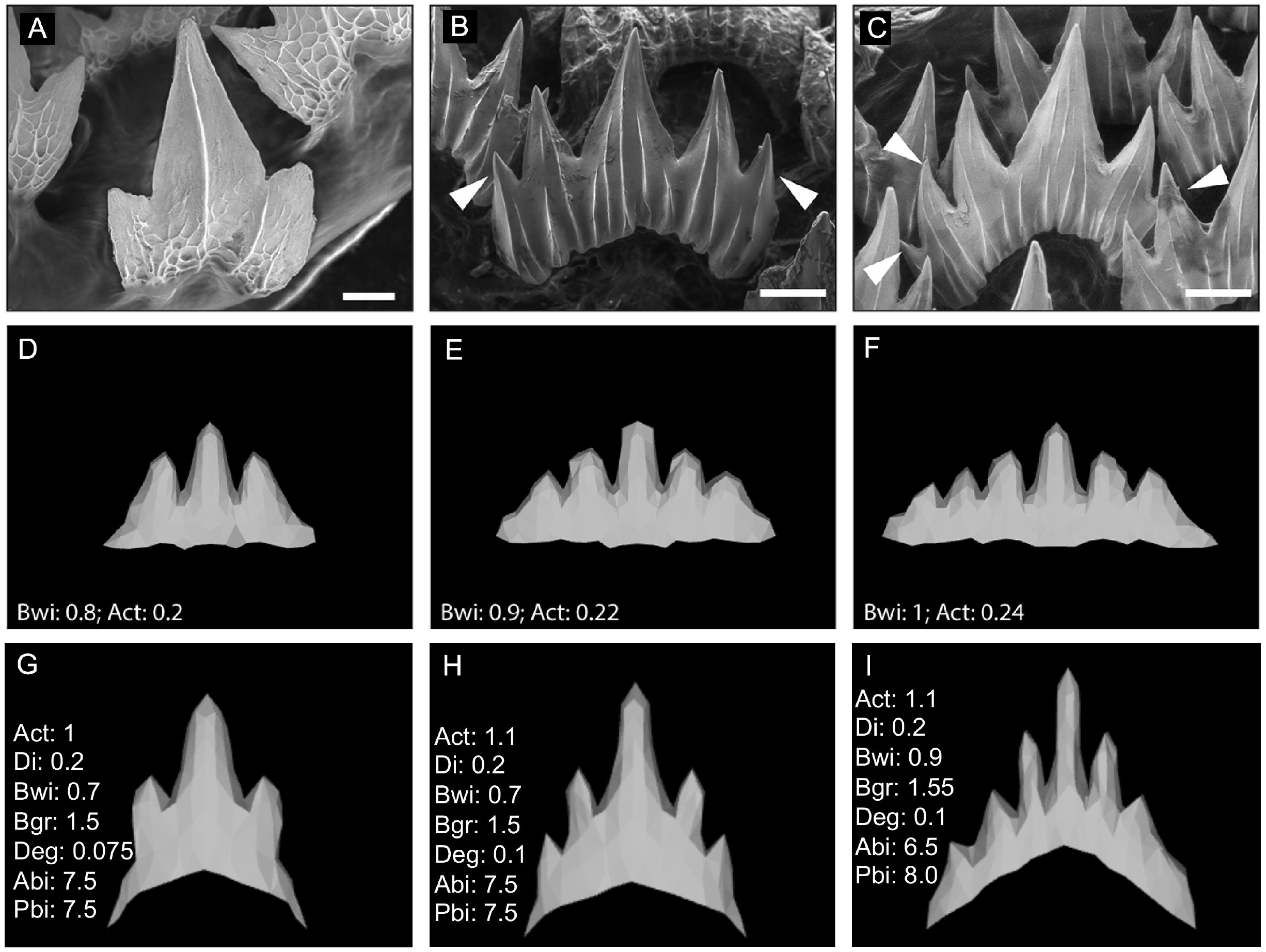
*In silico* modelling of the catshark dentition. Wild type SEM images of embryonic (A) and juvenile (B and C) catshark samples show a shift in cusp number from 3 to 6/7 cusps during ontogeny. The computational model ‘ToothMaker’ [5] was used to generate *in silico* models of the dentition (D-F; J-O). 11000 iterations of the model were run when modelling dentition, parameters for the models are available in the supplementary information (Table S1 and S2). Adjusting parameters reflecting underlying genetic networks, including Wnt signalling and the physical starting size shift teeth from a tricuspid (D) to a 5 (E) or 6 (F) or 7 cusped dentition. White arrowheads represent the addition of extra cusps during ontogeny. Scale bars are 50μm, and 200μm in B and C.

In contrast, upregulation of canonical Wnt signalling via CHIR99021 treatment resulted in teeth more similar in appearance to the standard catshark dentition. Lateral CHIR99021 teeth are clearly larger and wider than controls (Figure 7A and B), with regions of the teeth appearing to initiate the formation of ectopic cusps, although these do not always fully form (Figure 7B: white arrowheads). Interestingly, within other regions of the jaw we do see the development of fully formed 4th cusps (Figure 8C2 and C3). This is similar to the phenotypes we see in juvenile catsharks, which develop teeth with up to 7 cusps. The development of 4th cusps can also be seen within some of the control teeth (Figure 8A3). However, the proportion of teeth exhibiting 4th cusps is significantly lower in control samples (1%), than CHIR99021 treated samples (12%) (chi-square: X-squared = 16.887, df = 1, p-value <0.001). Furthermore, the position and size of cusps in DMSO treated samples is very consistent. There is a clear single primary cusp, with two secondary cusps equal in size. Extra cusps develop laterally and are smaller in size than the initial secondary cusps. This is not the case in CHIR99021 treated specimens. We note cases whereby secondary cusps are enlarged and can be difficult to distinguish apart from the primary cusp. We also observe teeth bearing multiple small cusps which resemble serrated teeth. The defects observed in final tooth shape highlight an important role for Wnt during dental morphogenesis. The specific directional shifts in the number of cusps associated with down and up regulation through IWR-1-endo and CHIR99021 treatment respectively, implicate canonical Wnt in the regulation of cusp development and therefore possibly in EK signalling.

### In silico modelling of tooth shape following perturbation of Wnt signalling

Salazar-Ciudad & Jernvall [5] developed a computational model (ToothMaker) capable of modelling vertebrate dentitions based on 12 cellular and 14 gene-network parameters with predetermined values. In order to establish the baseline parameters capable of generating a wild-type catshark tooth, we started with parameters established in modelling the seal dentition [5]. We iterated through each parameter at 10% intervals; using a process of elimination to refine the final tooth parameters (Figure 8; Figure S4). A shift in cusp number is observed throughout ontogeny, with an increase from 3 cusps in the embryo, to 5 or 6 cusps in juveniles (Fig 8, A-C). Simultaneously changing the width of the initial tooth site (Bwi) together with activation (Act) is capable of recreating 5 and 6 cusped teeth (Fig 8, D-F), as seen in the baseline seal model simulations. As sharks grow, the size of new dental generations increases (Fig 8A-C). It is likely that an increase in the size of the initial tooth site generates larger teeth with more numerous cusps.

Following shifts in dental morphology as a result of canonical Wnt manipulation, we sought to determine which parameters were capable of generating comparable shifts *in silico* (Figure 9). We identified two genetic parameters, which were individually capable of generating comparable phenotypes to the chemically treated samples. Both decreasing the activator auto-activation (Act = 0.1) and increasing the inhibition of the activator (Inh = 8) (Figure 9), resulted in unicuspid phenotypes strikingly similar to IWR-1-endo treated samples (Fig 9G). These results are biologically relevant. IWR-1-endo increases the production of the canonical Wnt inhibitor Axin2 [39]. The resulting function of increasing Axin2 *in vivo*, can be equally compared to increasing the inhibitor (Inh) or decreasing the effect of an activator (Act) as a result of increased inhibition, *in silico*. The converse is true for CHIR99021, which stabilises cytoplasmic β-catenin leading to upregulated canonical Wnt signalling [41]. This can be compared to decreasing the effect of an inhibitor, which would lead to higher cytoplasmic β-catenin (Inh = 1.2) or increasing the activator itself (Act = 0.22) (Fig 9C).

**Figure 9.**
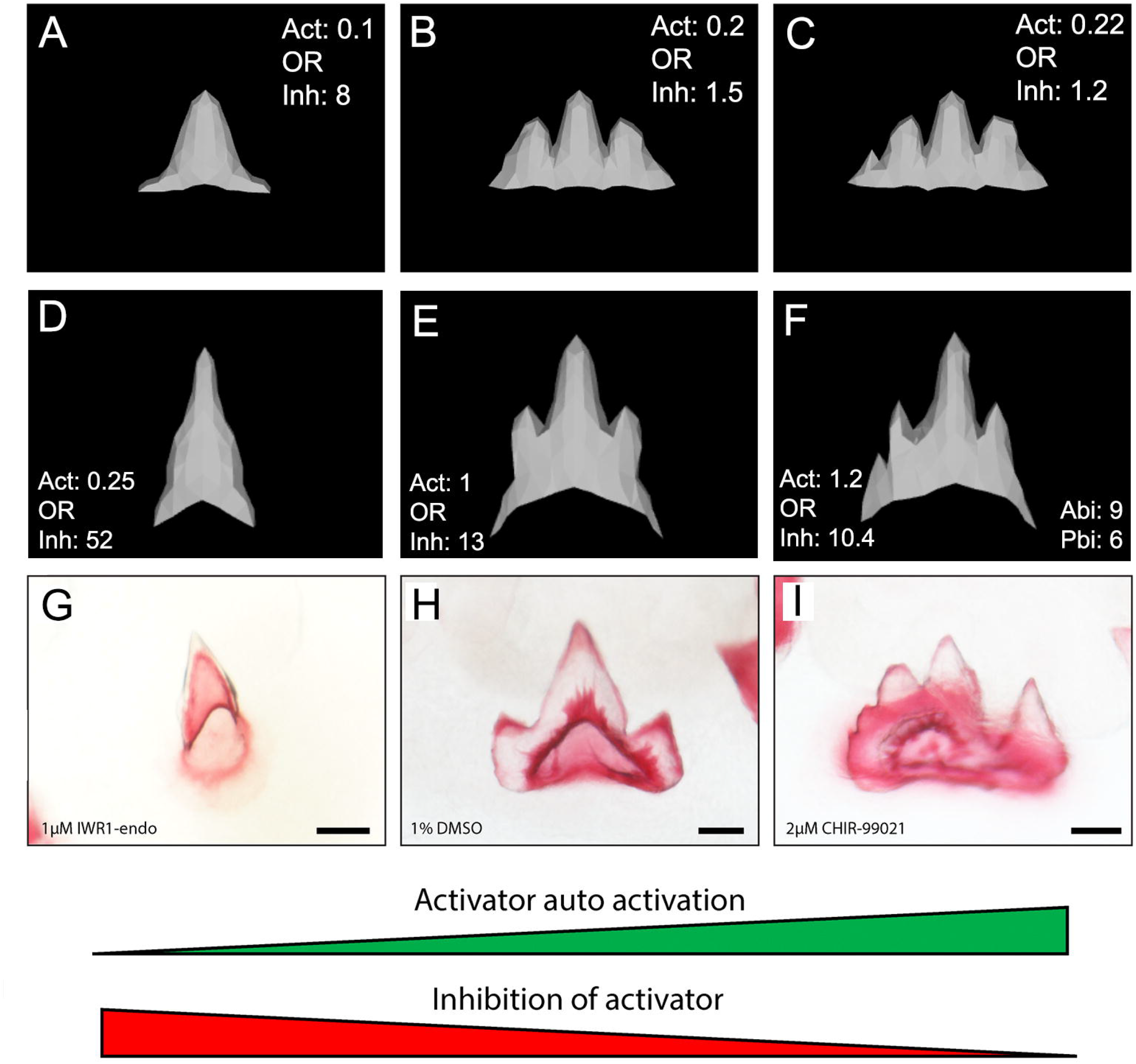
*In silico* modelling recreates catshark tooth phenotypes comparable to canonical Wnt manipulation. *in silico* modelling (A-C) is also capable of reproducing dental morphologies observed following chemical treatment (D-F) compared to the base tooth model from the simulation ToothMaker (seal tooth phenotype; G-L). 1μM IWR-1-endo resulted in unicuspid teeth (A, D), whereas 2μM CHIR9902 (C, F) resulted in the development of supernumerary cusps. Either an increase in activator auto activation (Act) (G-I) or decrease in inhibition of activator (Inh) (J-L), is sufficient to shift teeth from a unicuspid (A, D, G, J) to tricuspid (B, H, K) and quadricuspid (C, F, I, L) morphology. 11000 iterations of the model were run when modelling the effect of small molecule treatment. Scale bars are 50μm in D-F.

The phenotypes generated by the model resulted in both wider teeth, and an increase in cusp number (Figure 8). This is directly comparable to the phenotypes observed following treatment with CHIR99021 (Figure 9F and I). In the model, the regulation of the inhibitor and activator stems from the EK signalling centre [5]. The recreation of comparable phenotypes through the alteration of biologically relevant genetic parameters supports our findings, which demonstrate a critical role of canonical Wnt signalling in the regulation of cusp number from the catshark EK (Figure 8 and 9). Our findings show that increasing canonical Wnt activation results in teeth more similar to the adult phenotype, seen through an increase in cusp number following CHIR99021 treatment (Fig 9).

After modelling a wild-type tricuspid tooth in correspondence with the software developers (J. Jernvall, *pers*. *comm*.), we iterated through the parameters to recreate the effects of the drug treatment *in silico* (Fig.9). The 4-cusp-offset phenotype observed with the CHIR99021 Wnt activator was replicated in the model by increasing the auto-activator parameter by 20%, combined with the introduction of an anterior-posterior bias (Figure 9F, I; Supp. Table S1). This bias may result from accessibility of the drug through the tissue. The one cusp phenotype following IWR-1-endo Wnt-inhibitor treatment was replicated by increasing the inhibition of activator four-fold (Figure 9D, G; Supp. Table S1). Of note, increasing the autoactivation parameter had the same effect as decreasing the inhibitor parameter by an equivalent amount. For example, decreasing the activator-autoactivation to a quarter gives the same effect as increasing the inhibitor value four-fold. The same phenotype could also be achieved by the combined action of decreasing the activator by half and increasing the inhibitor two-fold.

When attempting to replicate the normal tooth development in ontogeny from tri-cuspid through to the 7-cusp phenotype, we found that there were two parameter-modules which influenced the final shape (Seal-type: Figure 8D-F; and shark-type: Figure 8G-I; Supplemental Table S2). The first module concerns the balance of the activatory and inhibitory propensity of the gene-network. Starting from a tricuspid baseline (Figure 8G), a 10%-20% increase in activator-autoactivation (or decrease in inhibition) results in a 5-cusp tooth (Figure 8H). Increasing another factor, the protein degradation by 33% gives the same result, and a combination of the two gives a sharper 5-cusps (Figure 8E and H; Supp. Table S2). Both of these properties could be reflective of changes to underlying Wnt signalling, with increased activator/decreased inhibitor directly, or a faster turnover of components favouring activation. Protein degradation is a key component of Wnt signalling activity, of which the GSK3β complex which degrades β-catenin is just one example [45,46]. Both the adjustment to protein degradation and activator/inhibitor levels needs to be met to move toward the 7-cusp phenotype (Figure 8I), and a concordant increase in inhibitor diffusion. Also required are adjustments to the second module of parameters, which concerns physical characteristics. An increase to the original border width of the tooth site and the border-growth rate is necessary to accommodate the larger tooth (Figure 8I, Supp. Table S2). In *S. canicula*, tooth buds are closely packed and during development form an inhibitory zone between units, which could represent a constraint on border size. Thus, the shift from 5 to 7-cusps is more complex than increasing the overall level of Wnt activation, simply doing so results in additional enamel knots forming outside of the anterior-posterior axis. These intricacies over shape control are perhaps enabled by the complexity of Wnt signalling, the number of interacting molecules permitting a wider range of phenotypes. To obtain 6-cusp teeth, the anterior-posterior bias can be skewed. Combined with the wider initial border, in this case one larger cusp still sits centrally, as seen in wildtype shark teeth (Figure 8A-C).

## Discussion

Overall, our results demonstrate that an enamel knot (EK)-like signalling centre regulates the formation of dental cusps in the catshark, with canonical Wnt signalling playing an important role in this process. We identify restricted expression of Fgf markers within the non-proliferative apical dental epithelium corresponding to the EK in mammals, and highlight shifts in cusp number and tooth shape following canonical Wnt manipulation, with similar results simulated via *in silico* modelling.

### Sharks possess an enamel knot-like signalling centre

Low levels of proliferation at the apical tip of the developing tooth have hinted at the presence of an EK in sharks [21], although three-dimensional restriction of signalling molecules within a distinct signalling centre is yet to be described. We note the expression of *fgf10* and the canonical Wnt inhibitor *dkk1* within the non-proliferative dental epithelium (Fig 3). Furthermore, our whole mount in situ hybridisation data reveals restricted upregulation of *fgf3* and *fgf10* within both primary and secondary cusp regions (Fig 4) and therefore the presence of a spatially restricted dental epithelial signalling centre. In mammals, the expression of EK markers precedes dental shape change [37]. Similarly, we observe the restricted expression of *fgf3* (Fig 2F) and *fgf10* (Fig 3A) very early during dental morphogenesis, prior to the establishment of the overall tooth shape. This suggests that signalling precedes dental shape change and suggests that Fgf signalling may be driving this process as in mammals [6,32]. Further investigation of EK markers prior to dental morphogenesis is key in discerning the role and regulation of EK driven dental morphogenesis in the shark.

Importantly, the mammalian EK itself is unable to respond to the Fgf signals, which it emits as it lacks Fgf receptors [47]. Instead, these receptors are found within adjacent dental epithelial cells, and are important for the induction of differential proliferation between the EK and surrounding epithelial tissue [32]. Further research is needed to identify whether this lack of receptors predates the evolution of the mammalian dentition, and whether it is fundamentally required for the formation of cusped teeth. Given the conservation of localised Fgf signalling within the shark EK, we hypothesise that a similar lack of receptors drives differential proliferation between the EK and surrounding dental epithelium in the basal gnathostome lineage.

Various developmental characteristics have been used to define the presence of an EK, including spatial restriction of the signalling centre, a lack of cell proliferation, and apoptosis of the epithelial cells within the EK. Apoptosis has been implicated as an important developmental process in regulating the silencing of embryonic signalling centres [32,37]. Apoptosis has been described in both primary and secondary mammalian EK through TUNEL assays and the expression of the pre-apoptotic marker, p21 [10,37]. Such assays have so far revealed a lack of apoptosis within the inner dental epithelium of reptiles [15,24] and the catshark [23] - this has been used to refute the presence of an EK altogether [23]. However, it has also been shown that mice develop cusps following the loss of apoptosis in caspase-3 deficient mice [48]. Although morphological defects are identified in non-apoptotic mouse molars [48], these results suggest that apoptosis is not fundamentally required in the formation of dental cusps [19].

Whilst there are observable differences in signalling between the mammalian and chondrichthyan EKs [23], this does not refute the presence of an EK in sharks. Instead, these differences highlight potential lineage specific modifications which have led to the diversification of vertebrate dental morphology. Given the presence of enameloid, as opposed to ‘true’ enamel within the Chondrichthyes [49], it has been proposed to term this signalling centre in sharks the ‘enameloid knot’ [21].

Our results reveal a complete lack of *bmp4* expression within the non-proliferative apical dental epithelium. *bmp4* expression in surrounding tissues suggests that unlike in mammals, where Bmp4 is secondarily upregulated within the EK, bmp4 may play a role in restricting the expression of other signalling molecules to the EK in the shark. Furthermore, shh is a key marker of the EK in mammals. Although we note its expression within the apical dental epithelium, there is little downregulation of its expression within the inter-cusp dental epithelium. Asymmetric *shh* expression is thought to regulate polarised growth of the developing feather bud [50] as it does in the zone of polarising activity (ZPA) within the vertebrate limb bud [51]. It is possible that shh is playing a similar role in teeth of sharks; shh may be regulating polarised growth of the tooth along the oral/aboral axis, but not the medial/lateral axis along which the cusps form. Despite differences in the gene specific expression patterns of the mammalian and chondrichthyan EKs, representative markers of canonical Wnt, Fgf, Bmp and Hh signalling are all found expressed within the apical dental epithelium.

### Canonical Wnt signalling as a regulator of natural shark dental variation

Chondrichthyan dental morphology is highly variable between species [25,52]. This variability allows for fossil samples to be commonly identified on their dentition alone [53]. Variation in chondrichthyan tooth shape is typically observable as a modification of the cusp. Examples include: the multi-cuspid saw-like teeth of the sixgill shark (*Hexanchus griseus*); the elongated primary cusp of mako shark teeth (*Isurus oxyrinchus*); serrated teeth of the tiger shark (*Galeocerdo cuvier*); and the ontogenetic shift from tricuspid-heptacuspid teeth of the small-spotted catshark (*S. canicula*) [25,52]. Our small-molecule manipulations of the canonical Wnt signalling pathway result in a shift in cusp number and an increase in the variability of tooth shape. Furthermore, the diversity of phenotypes observed following treatment include: unicuspid, multicuspid and serrated-like teeth. Interestingly, the factors and developmental mechanisms that can lead to the formation of serrated teeth in any vertebrate are largely unknown.

Canonical Wnt pathway signalling is known to lie upstream of EK signalling during mammalian molar morphogenesis, with its upregulation and downregulation leading to supernumerary EKs and the formation of blunted cusps respectively [13,54]. In the shark, our findings indicate a similar role for Wnt signalling during dental morphogenesis. Here, Wnt manipulation dramatically disrupts cusp development. As a result, we speculate that alterations to the canonical Wnt pathway may also underlie the natural diversity of tooth shape found between chondrichthyan species. However, although our chemical manipulation phenotypes match what would be expected as a result of disrupted EK signalling, we cannot exclude the possibility that the observed shifts in shape arise as a result of changes in size of the teeth or developmental arrest of tooth morphogenesis. Analysis of canonical Wnt signalling targets within the EK following chemical manipulation, could provide further clarity as to which of these developmental processes is driving shape change.

An *in silico* model of dental morphogenesis developed by Salazar-Ciudad and Jernvall [5] demonstrates how dental shape can be regulated by a variety of cellular and genetic parameters in the EK. Subtle changes to these parameters lead to a drastic change in dental shape and cusp number. We show that small changes to two genetic parameters regulating activation (*Act*) or inhibition (*Inh*) of the activator in the EK, is sufficient in generating teeth representative of upregulation and downregulation of canonical Wnt signalling, respectively. However, the parameters necessary to produce an ontogenetic shift in shark-specific tooth shape may be more complex (Figure 9). The similarities observed following Wnt treatments and *in silico* EK signalling manipulations, provide both an element of validation for the *in silico* model and further evidence of a role for canonical Wnt signalling in the chondrichthyan EK.

### Was an EK-like signalling centre present prior to the evolution of teeth?

Odontodes (tooth-like structures) are thought to have initially evolved outside of the oral cavity, with teeth arising through co-option of the underlying odontode gene regulatory network [27,55,56]. Dermal denticles (non-regenerative odontodes present on the skin surface of chondrichthyans) and teeth are deeply homologous, sharing a high degree of structural and developmental conservation [27,57,58]. We observe identical expression patterns for *fgf3*, *bmp4* and *shh*, within and around the apical tip of developing teeth (Figure 2) and dermal denticles in the catshark [57,58]. As a result, we believe it is likely that the origin of an apical epithelial signalling centre, regulating differential epithelial proliferation in epithelial appendages evolved early within the vertebrate lineage and likely predates the evolution of teeth. Therefore, to account for this regulation of shape among disparate developmental structures we propose a new term to reflect the conservation of this signalling unit, the ‘<u>Apical Epithelial Knot’</u> (AEK). This then suggests that this signalling centre is not limited to teeth but can be present in a number of protrusible developmental elements, including non-oral odontodes. The enamel knot term can be used exclusively for mammalian tooth signalling, due to the distinction between the enamel and enameloid capping mineral layer observed in fish teeth and denticles, and other odontodes.

## Conclusion

Although there are differences in the expression of key mammalian EK markers in the shark, there is also conservation of key Wnt, Fgf and Shh markers. Contrary to prior assertions, these results provide evidence of an EK signalling centre in an early vertebrate lineage. We therefore suggest that mammalian-specific gene expression patterns within the EK are merely lineage specific modifications of an ancestral signalling centre (AEK) that likely evolved prior to the appearance of oral teeth.

## Methods

### Animal husbandry

The University of Sheffield is a licensed establishment under the Animals (Scientific Procedures) Act 1986. All animals were culled by approved methods cited under Schedule 1 to the Act. Small-spotted catshark embryos (*S. canicula*) were obtained from North Wales Biologicals, Bangor, UK. Embryos were raised in recirculating artificial seawater (Instant Ocean) at 16°C. At the required stage, embryos were anaesthetised using 300mg/L MS-222 and fixed overnight in 4% paraformaldehyde at 4°C. Samples were then dehydrated through a graded series of DEPC-PBS/EtOH and kept at −20°C.

### Sectioning and Histology

Following dehydration, samples were cleared with xylene and embedded in paraffin. 14μm sagittal sections were obtained using a Leica RM2145 microtome. For histological study, sections were stained with 50% Haematoxylin Gill no.3 and Eosin Y. Slides were mounted with Fluoromount (Sigma) and imaged using a BX51 Olympus compound microscope.

### Scanning Electron Microscopy (SEM)

SEM images were obtained using a Hitachi TM3030Plus Benchtop SEM at 15000 Volts.

### In situ *hybridisation probe synthesis*

Protein coding sequences for S. canicula were obtained from a de-novo transcriptome assembly (Thiery et al. *unpublished*). Sequences were compared with a range of other vertebrate sequences taken from ensembl.org in order to verify sequence identity. *S. canicula* total RNA was extracted using phenol/chloroform phase separation and cleaned through EtOH/LiCL precipitation. RT-cDNA was made using the RETRO script 1710 kit (Ambion). Probes were made using forward and reverse primers designed through Primer3. Primer sequences are available in the supplementary information (Figure S5). Probes were chosen to be ~400-800bp in length. Sequences of interest were amplified from the cDNA through PCR and ligated into the pGEM-T-Easy vector (Promega). Ligation products were cloned into JM109 cells. Plasmid DNA was then extracted from chosen colonies using a Qiaprep spin Mini-prep kit (Qiagen) and sequenced (Applied Biosystems’ 3730 DNA Analyser) through the Core Genomics Facility, University of Sheffield. Verified vectors were then amplified through PCR and used as a template for probe synthesis. Sense and anti-sense probes were made using a Riboprobe Systems kit (Promega) and SP6/T7 polymerases (Promega). Probes were labelled with Digoxigenin-11-UTP (Roche) for detection during in situ hybridisation. A final EtOH precipitation step was carried out to purify the RNA probe.

### *Section* in situ *hybridisation*

Sagittal paraffin sections were obtained as previously described. Slides were deparaffinised using Xylene and rehydrated through a graded series of EtOH/PBS. Slides were incubated in pre-heated pre-hybridisation solution pH 6 [250ml deionised-formamide, 125ml 20x saline sodium citrate (SSC), 5ml 1M sodium citrate, 500μl Tween-20 and 119.7ml DEPC-treated ddH20] at 61°C for 2 hours. Slides were transferred to pre-heated pre-hybridisation solution containing DIG labelled RNA probe (1:500) and incubated overnight at 61°C. The following day, slides underwent a series of 61°C SSC stringency washes to remove unspecific probe binding [2×30m 50:50 pre-hybridisation solution:2x SSC; 2×30m 2x SSC; 2×30m 0.2x SSC]. Following the stringency washes, samples were incubated in blocking solution (2% Roche Blocking Reagent (Roche)) for 2hr at room temperature and then incubated in blocking solution containing anti-Digoxigenin-AP antibody (1:2000; Roche) overnight at 4°C. Excess antibody was washed off through 6×1hr MAB-T (0.1% tween-20) washes. Slides were then washed in NTMT and colour reacted with BM-purple (Roche) at room temperature and left until sufficient colouration had taken place. Following the colour reaction, a DAPI nuclear counterstain (1μg/ml) was carried out before mounting the slides using Fluoromount (Sigma). Images were taken using a BX51 Olympus compound microscope. Images were contrast enhanced and merged in Adobe Photoshop.

### *Double* in situ *hybridisation/immunohistochemistry*

For double in situ hybridisation/immunohistochemistry, samples first underwent in situ hybridisation as previously described. Immediately after colour reaction, samples were fixed for 1 minute in 4% paraformaldehyde in PBS. Samples were then blocked with 5% goat serum and 1% bovine serum albumin in PBS-T (0.05% tween-20). Blocking solution was replaced with blocking solution containing mouse anti-PCNA primary antibody (ab29; Abcam) at a concentration of 1:2000. Goat anti-rabbit Alexa-Fluor 647 (1:250) (A-20721245; Thermo) and goat anti-mouse Alexa-Fluor 488 (1:250) (A-11-001; Thermo) secondary antibodies were used for immunodetection. Samples were counterstained with DAPI (1μg/ml) and mounted using Fluoromount (Sigma). Images were taken using a BX51 Olympus compound microscope. Images were contrast enhanced and merged in Adobe Photoshop.

### *Whole mount* in situ *hybridisation*

Whole mount in situ hybridisation was carried out in accordance with the section in situ hybridisation protocol with some minor modifications. Following rehydration, samples were treated with 0.2μg/ml proteinase K for 1hr at room temperature and then fixed for 20m in 4% paraformaldehyde in PBS. Samples were then placed in pre-hybridisation and probe solution as previously described. Stringency washes were carried out at 61°C [3×30m 2xSSC-T (0.05% tween-20); 3×30m 0.2xSSC-T (0.05% tween-20)]. Blocking, antibody incubation and colour reaction were carried out as previously described. Following colour reaction, samples were stored in PBS with 10% EtOH.

### Small molecule Wnt perturbations

10mM IWR-1-endo (product no) and 5mM CHIR99021 (product no) stock solutions were made using dimethyl sulfoxide (DMSO) as a solvent. At ~100dpf (mid stage 32), catshark samples were extracted from their egg cases and incubated in 70ml polypropylene containers. Samples were treated with 1μM (1:10000 stock dilution) IWR-1-endo (N = 8), 2μM (1:2500 stock dilution) CHIR99021 (N = 8). 0.1% DMSO was used as a control (N = 7). Chemical stock solutions were diluted in artificial seawater with 1% Penicillin/Streptomycin. Samples were treated with 20ml of solution, which was replaced every two days for a total treatment period of two weeks. Following treatment, samples were raised in artificial seawater for a further 4 weeks. After recovery, samples were sacrificed using 300mg/L MS-222 and fixed overnight in 4% paraformaldehyde at 4°C. Samples were then stained with 0.02% alizarin red in 0.1% KOH overnight in the dark and subsequently cleared in 0.1% KOH. Once residual alizarin red had been removed, samples were transferred into glycerol through a glycerol/0.1%KOH graded series and imaged using a Nikon SMZ1500.

### Geometric morphometric analysis

Images taken from the small molecule treated clear and stained specimens were analysed using 2-D geometric morphometrics. Images of the three most lateral teeth on both the left and right side of the lower jaw were included in the analysis. TPS files containing the treatment images were generated in tpsUtil [59,60]. Landmark coordinates were assigned using tpsDig2 [59]. Two fixed landmarks were placed on each tooth, one at the tip of the primary cusp and one at the base of the tooth. 18 sliding-semi landmarks were then distributed evenly between these two points on either side of the tooth. TPS files were analysed using the R package geomorph [61]. A generalised Procrustes analysis (GPA) was carried out in order to rotate, centre and rescale image coordinates. Following GPA, we measured the effect of treatment and sample on Centroid size using a linear model, with both factors included as fixed effects. Comparisons between treatments were made using a post hoc Tukey test. Next, we assessed the contribution of treatment and sample to shape (Procrustes aligned coordinates) using a linear model procD.lm function in the geomorph R package [61] with both factors included as fixed effects. In order to plot the shape data, a principle component analysis was conducted on the Procrustes aligned co-ordinates and plotted based on the principle components which explain the most variation in the data (Figure 6). We measured the variance for principal component 1 within treatments and carried out a pairwise F-test to test for changes in variance as a result of treatment.

### ToothMaker modelling of catshark dentition

*In silico* models of the catshark dentition through ontogeny were generated using the computational model ToothMaker [5], together with the equivalent cusp number changes in the baseline seal tooth parameters [5]. Initial shark phenotype baseline parameters were established with assistance from the program creator (Jukka Jernvall, *Pers*. *Comm*.). We iterated through each parameter at 10% intervals; using a process of elimination to refine the final wild type tooth parameters for the modified seal tooth phenotypes to so cusp number equivalence to the catshark (Figure S4). Trial and error was then used to determine which parameters were capable of generating teeth resembling those produced following small molecule treatment (Supp. Table S1) and in normal development (Supp. Table S2). 11000 iterations of the model were run when modelling the ontogenetic shifts in the dentition.

## Acknowledgements

We are grateful to Jukka Jernvall for his invaluable insights and advice with regard to the Tooth-Maker simulation software. We thank Keith Hunter and Abigail Tucker for useful comments on previous versions of this manuscript. Funding: University of Sheffield Adapting to the Challenges of Changing Environment Doctoral Training Programme (ACCE) (to RLC and APT); Natural Environment Research Council grant NE/K014595/1 (to GJF); Leverhulme Trust grant RPG-211 (to GJF).

## References

1. Jernvall J, Thesleff I. 2012 Tooth shape formation and tooth renewal: evolving with the same signals. Development 139, 3487–3497.

2. Tucker AS, Fraser GJ. 2014 Evolution and developmental diversity of tooth regeneration. Semin. Cell Dev. Biol. 25-26, 71–80.

3. Zhang YD, Chen Z, Song YQ, Chao LIU, Chen YP. 2005 Making a tooth: growth factors, transcription factors, and stem cells. Cell Research. 15, 301–316. (doi:10.1038/sj.cr.7290299)

4. Thesleff I, Sharpe P. 1997 Signalling networks regulating dental development. Mech. Dev. 67, 111–123.

5. Salazar-Ciudad I, Jernvall J. 2010 A computational model of teeth and the developmental origins of morphological variation. Nature 464, 583–586.

6. Jernvall J, Kettunen P, Karavanova I, Martin LB, Thesleff I. 1994 Evidence for the role of the enamel knot as a control center in mammalian tooth cusp formation: non-dividing cells express growth stimulating Fgf-4 gene. Int. J. Dev. Biol. 38, 463–469.

7. Sumigray KD, Terwilliger M, Lechler T. 2018 Morphogenesis and Compartmentalization of the Intestinal Crypt. Dev. Cell 45, 183–197.e5.

8. Ahtiainen L, Lefebvre S, Lindfors PH, Renvoisé E, Shirokova V, Vartiainen MK, Thesleff I, Mikkola ML. 2014 Directional cell migration, but not proliferation, drives hair placode morphogenesis. Dev. Cell 28, 588–602.

9. Mustonen T, Tümmers M, Mikami T, Itoh N, Zhang N, Gridley T, Thesleff I. 2002 Lunatic fringe, FGF, and BMP regulate the Notch pathway during epithelial morphogenesis of teeth. Dev. Biol. 248, 281–293.

10. Jernvall J, Aberg T, Kettunen P, Keränen S, Thesleff I. 1998 The life history of an embryonic signaling center: BMP-4 induces p21 and is associated with apoptosis in the mouse tooth enamel knot. Development 125, 161–169.

11. Salazar-Ciudad I. 2012 Tooth patterning and evolution. Curr. Opin. Genet. Dev. 22, 585–592.

12. Kratochwil K, Galceran J, Tontsch S, Roth W, Grosschedl R. 2002 FGF4, a direct target of LEF1 and Wnt signaling, can rescue the arrest of tooth organogenesis in Lef1(-/-) mice. Genes Dev. 16, 3173–3185.

13. Liu F et al. 2008 Wnt/beta-catenin signaling directs multiple stages of tooth morphogenesis. Dev. Biol. 313, 210–224.

14. Jia S, Kwon H-JE, Lan Y, Zhou J, Liu H, Jiang R. 2016 Bmp4-Msx1 signaling and Osr2 control tooth organogenesis through antagonistic regulation of secreted Wnt antagonists. Dev. Biol. 420, 110–119.

15. Buchtová M, Handrigan GR, Tucker AS, Lozanoff S, Town L, Fu K, Diewert VM, Wicking C, Richman JM. 2008 Initiation and patterning of the snake dentition are dependent on Sonic hedgehog signaling. Dev. Biol. 319, 132–145.

16. Weeks O, Bhullar B-AS, Abzhanov A. 2013 Molecular characterization of dental development in a toothed archosaur, the American alligator Alligator mississippiensis. Evol. Dev. 15, 393–405.

17. Zahradnicek O, Buchtova M, Dosedelova H, Tucker AS. 2014 The development of complex tooth shape in reptiles. Front. Physiol. 5, 74.

18. Handrigan GR, Richman JM. 2011 Unicuspid and bicuspid tooth crown formation in squamates. J. Exp. Zool. B Mol. Dev. Evol. 316, 598–608.

19. Richman JM, Handrigan GR. 2011 Reptilian tooth development. Genesis 49, 247–260.

20. Fraser GJ, Bloomquist RF, Streelman JT. 2013 Common developmental pathways link tooth shape to regeneration. Dev. Biol. 377, 399–414.

21. Rasch LJ, Martin KJ, Cooper RL, Metscher BD, Underwood CJ, Fraser GJ. 2016 An ancient dental gene set governs development and continuous regeneration of teeth in sharks. Dev. Biol. 415, 347–370.

22. Rasch LJ, Cooper RL, Underwood C, Dillard WA, Thiery AP, Fraser GJ. 2020 Development and regeneration of the crushing dentition in skates (Rajidae). Dev. Biol. 466, 59–72.

23. Debiais-Thibaud M, Chiori R, Enault S, Oulion S, Germon I, Martinand-Mari C, Casane D, Borday-Birraux V. 2015 Tooth and scale morphogenesis in shark: an alternative process to the mammalian enamel knot system. BMC Evol. Biol. 15, 292.

24. Handrigan GR, Richman JM. 2010 Autocrine and paracrine Shh signaling are necessary for tooth morphogenesis, but not tooth replacement in snakes and lizards (Squamata). Dev. Biol. 337, 171–186.

25. Jambura PL, Türtscher J, Kindlimann R, Metscher B, Pfaff C, Stumpf S, Weber GW, Kriwet J. 2020 Evolutionary trajectories of tooth histology patterns in modern sharks (Chondrichthyes, Elasmobranchii). J. Anat. 236, 753–771.

26. Fraser GJ, Graham A, Smith MM. 2004 Conserved deployment of genes during odontogenesis across osteichthyans. Proc. Biol. Sci. 271, 2311–2317.

27. Martin KJ, Rasch LJ, Cooper RL, Metscher BD, Johanson Z, Fraser GJ. 2016 Sox2+ progenitors in sharks link taste development with the evolution of regenerative teeth from denticles. Proc. Natl. Acad. Sci. U. S. A. 113, 14769–14774.

28. Thiery AP, Shono T, Kurokawa D, Britz R, Johanson Z, Fraser GJ. 2017 Spatially restricted dental regeneration drives pufferfish beak development. Proc. Natl. Acad. Sci. U. S. A. 114, E4425–E4434.

29. Ballard WW, Mellinger J, Lechenault H. 1993 A series of normal stages for development ofScyliorhinus canicula, the lesser spotted dogfish(Chondrichthyes: Scyliorhinidae). Journal of Experimental Zoology. 267, 318–336. (doi:10.1002/jez.1402670309)

30. Wu X, Li Y, Wang F, Hu L, Li Y, Wang J, Zhang C, Wang S. 2017 Spatiotemporal Expression of Wnt/β-catenin Signaling during Morphogenesis and Odontogenesis of Deciduous Molar in Miniature Pig. Int. J. Biol. Sci. 13, 1082–1091.

31. Sarkar L, Sharpe PT. 1999 Expression of Wnt signalling pathway genes during tooth development. Mech. Dev. 85, 197–200.

32. Kettunen P, Laurikkala J, Itäranta P, Vainio S, Itoh N, Thesleff I. 2000 Associations of FGF-3 and FGF-10 with signaling networks regulating tooth morphogenesis. Dev. Dyn. 219, 322–332.

33. Moustakas JE, Smith KK, Hlusko LJ. 2011 Evolution and development of the mammalian dentition: insights from the marsupial Monodelphis domestica. Dev. Dyn. 240, 232–239.

34. Vainio S, Karavanova I, Jowett A, Thesleff I. 1993 Identification of BMP-4 as a signal mediating secondary induction between epithelial and mesenchymal tissues during early tooth development. Cell 75, 45–58.

35. Massagué J. 2012 TGFβ signalling in context. Nat. Rev. Mol. Cell Biol. 13, 616–630.

36. Xu X, Jeong L, Han J, Ito Y, Bringas P Jr, Chai Y. 2003 Developmental expression of Smad1-7 suggests critical function of TGF-beta/BMP signaling in regulating epithelial-mesenchymal interaction during tooth morphogenesis. Int. J. Dev. Biol. 47, 31–39.

37. Vaahtokari A, Åberg T, Jernvall J, Keränen S, Thesleff I. 1996 The enamel knot as a signaling center in the developing mouse tooth. Mechanisms of Development. 54, 39–43. (doi:10.1016/0925-4773(95)00459-9)

38. Hardcastle Z, Mo R, Hui CC, Sharpe PT. 1998 The Shh signalling pathway in tooth development: defects in Gli2 and Gli3 mutants. Development 125, 2803–2811.

39. Chen B et al. 2009 Small molecule-mediated disruption of Wnt-dependent signaling in tissue regeneration and cancer. Nat. Chem. Biol. 5, 100–107.

40. Ring DB et al. 2003 Selective glycogen synthase kinase 3 inhibitors potentiate insulin activation of glucose transport and utilization in vitro and in vivo. Diabetes 52, 588–595.

41. Wagman AS, Johnson KW, Bussiere DE. 2004 Discovery and development of GSK3 inhibitors for the treatment of type 2 diabetes. Curr. Pharm. Des. 10, 1105–1137.

42. Seetah K, Cucchi T, Dobney K, Barker G. 2014 A geometric morphometric reevaluation of the use of dental form to explore differences in horse (Equus caballus) populations and its potential zooarchaeological application. Journal of Archaeological Science. 41, 904–910. (doi:10.1016/j.jas.2013.10.022)

43. Klingenberg CP. 2016 Size, shape, and form: concepts of allometry in geometric morphometrics. Dev. Genes Evol. 226, 113–137.

44. Perez SI, Bernal V, Gonzalez PN. 2006 Differences between sliding semi-landmark methods in geometric morphometrics, with an application to human craniofacial and dental variation. J. Anat. 208, 769–784.

45. Gao C, Xiao G, Hu J. 2014 Regulation of Wnt/β-catenin signaling by posttranslational modifications. Cell & Bioscience. 4, 13. (doi:10.1186/2045-3701-4-13)

46. Tauriello DVF, Maurice MM. 2010 The various roles of ubiquitin in Wnt pathway regulation. Cell Cycle. 9, 3724–3733. (doi:10.4161/cc.9.18.13204)

47. Kettunen P, Karavanova I, Thesleff I. 1998 Responsiveness of developing dental tissues to fibroblast growth factors: expression of splicing alternatives of FGFR1, −2, −3, and of FGFR4; and stimulation of cell proliferation by FGF-2, −4, −8, and −9. Dev. Genet. 22, 374–385.

48. Matalova E, Sharpe PT, Lakhani SA, Roth KA, Flavell RA, Setkova J, Misek I, Tucker AS. 2006 Molar tooth development in caspase-3 deficient mice. Int. J. Dev. Biol. 50, 491–497.

49. Gillis JA, Donoghue PCJ. 2007 The homology and phylogeny of chondrichthyan tooth enameloid. J. Morphol. 268, 33–49.

50. Ting-Berreth SA, Chuong CM. 1996 Sonic Hedgehog in feather morphogenesis: induction of mesenchymal condensation and association with cell death. Dev. Dyn. 207, 157–170.

51. Riddle RD, Johnson RL, Laufer E, Tabin C. 1993 Sonic hedgehog mediates the polarizing activity of the ZPA. Cell 75, 1401–1416.

52. Corn KA, Farina SC, Brash J, Summers AP. 2016 Modelling tooth–prey interactions in sharks: the importance of dynamic testing. Royal Society Open Science. 3, 160141. (doi:10.1098/rsos.160141)

53. † LBW, Whitenack † LB, Gottfried MD. 2010 A morphometric approach for addressing tooth-based species delimitation in fossil mako sharks,Isurus(Elasmobranchii: Lamniformes). Journal of Vertebrate Paleontology. 30, 17–25. (doi:10.1080/02724630903409055)

54. Järvinen E, Salazar-Ciudad I, Birchmeier W, Taketo MM, Jernvall J, Thesleff I. 2006 Continuous tooth generation in mouse is induced by activated epithelial Wnt/beta-catenin signaling. Proc. Natl. Acad. Sci. U. S. A. 103, 18627–18632.

55. Fraser GJ, Cerny R, Soukup V, Bronner-Fraser M, Streelman JT. 2010 The odontode explosion: the origin of tooth-like structures in vertebrates. Bioessays 32, 808–817.

56. Donoghue PCJ, Rücklin M. 2016 The ins and outs of the evolutionary origin of teeth. Evol. Dev. 18, 19–30.

57. Cooper RL, Martin KJ, Rasch LJ, Fraser GJ. 2017 Developing an ancient epithelial appendage: FGF signalling regulates early tail denticle formation in sharks. Evodevo 8, 8.

58. Cooper RL, Thiery AP, Fletcher AG, Delbarre DJ, Rasch LJ, Fraser GJ. 2018 An ancient Turing-like patterning mechanism regulates skin denticle development in sharks. Sci Adv 4, eaau5484.

59. Rohlf FJ. 2009 tpsDig. See http://life.bio.sunysb.edu/morph/soft-dataacq.html.

60. Rohlf JF. 2009 tpsUtil. See http://life.bio.sunysb.edu/morph/soft-utility.html.

61. Adams D Collyer M Kaliontzopoulou. 2018 Geomorph: Software for geometric morphometric analyses. R package version 3.0.6. See https://cran.r-project.org/package=geomorph.

