## Supplemental Table 1 for "An ancient dental signalling centre supports homology of an enamel knot in sharks"

| *Symbol* | *Parameter* | *1-Cusp* | ***3-Cusp*** | *4-Cusp* |
| --- | --- | --- | --- | --- |
| Egr | Epithelial Growth | 0.0225 | **0.0225** | 0.0225 |
| Mgr | Mesenchymal Growth | 200 | **200** | 200 |
| Rep | Cell Repulsion | 1 | **1** | 1 |
| Swi | Distance from 0 where borders defined | 0 | **0** | 0 |
| Adh | Cell Adhesion | 0.001 | **0.001** | 0.001 |
| Act | Activator Self-regulation | 1 | **1** | **1.2** |
| Inh | Inhibitor Strength | **52** | **13** | 13 |
| Sec | Secondary Signals | 0.03 | **0.03** | 0.03 |
| Da | Activator Diffusion | 0.2 | **0.2** | 0.2 |
| Di | Inhibitor Diffusion | 0.2 | **0.2** | 0.2 |
| Ds | Growth factor diffusion Rate | 0.2 | **0.2** | 0.2 |
| Int | Initial Inhibitor Threshold | 0.19 | **0.19** | 0.19 |
| Set | Secondary Signal Threshold | 0.95 | **0.95** | 0.95 |
| Boy | Mesenchymal Buoyancy | 0.1 | **0.1** | 0.1 |
| Dff | Differentiation Rate | 0.0004 | **0.0004** | 0.0004 |
| Bgr | Border Growth | 1.5 | **1.5** | 1.5 |
| Abi | Anterior Bias | 7.5 | **7.5** | 9 |
| Pbi | Posterior Bias | 7.5 | **7.5** | 6 |
| Bbi | Buccal Bias | 1 | **1** | 1 |
| Lbi | Lingual Bias | 1 | **1** | 1 |
| Rad | Radius of Initial Conditions | 2 | **2** | 2 |
| Deg | Protein Degradation Rate | 0.075 | **0.075** | 0.075 |
| Dgr | Downward Growth | 10000 | **10000** | 10000 |
| Ntr | Nucleus Traction | 0.00001 | **0.00001** | 0.00001 |
| Bwi | Border Width | 0.7 | **0.7** | 0.7 |
| Ina | Initial Activator Concentration | 0 | **0** | 0 |

**Supplementary Table S1.** ToothMaker parameters set for catshark tricuspid tooth, with modifications performed to replicate drug treatment phenotypes. (Salazar-Ciudad and Jernvall, 2010).
