## Supplemental Table 2 for "An ancient dental signalling centre supports homology of an enamel knot in sharks"

| *Symbol* | *Parameter* | ***3-cusp*** | *5-cusp* | *6-cusp* | *7-cusp* |
| --- | --- | --- | --- | --- | --- |
| Egr | Epithelial Growth | **0.0225** | 0.0225 | 0.0225 | 0.0225 |
| Mgr | Mesenchymal Growth | **200** | 200 | 200 | 200 |
| Rep | Cell Repulsion | **1** | 1 | 1 | 1 |
| Swi | Distance from 0 where borders defined | **0** | 0 | 0 | 0 |
| Adh | Cell Adhesion | **0.001** | 0.001 | 0.001 | 0.001 |
| Act | Activator Self-regulation | **1** | **1.1** | **1.1** | **1.1** |
| Inh | Inhibitor Strength | **13** | 13 | 13 | 13 |
| Sec | Secondary Signals | **0.03** | 0.03 | 0.03 | 0.03 |
| Da | Activator Diffusion | **0.2** | 0.2 | 0.2 | 0.2 |
| Di | Inhibitor Diffusion | **0.2** | 0.2 | **0.245** | **0.245** |
| Ds | Growth factor diffusion Rate | **0.2** | 0.2 | 0.2 | 0.2 |
| Int | Initial Inhibitor Threshold | **0.19** | 0.19 | 0.19 | 0.19 |
| Set | Secondary Signal Threshold | **0.95** | 0.95 | 0.95 | 0.95 |
| Boy | Mesenchymal Buoyancy | **0.1** | 0.1 | 0.1 | 0.1 |
| Dff | Differentiation Rate | **0.0004** | 0.0004 | 0.0004 | 0.0004 |
| Bgr | Border Growth | **1.5** | 1.5 | **1.55** | **1.55** |
| Abi | Anterior Bias | **7.5** | 7.5 | **6.5** | 7.5 |
| Pbi | Posterior Bias | **7.5** | 7.5 | **8.0** | 7.5 |
| Bbi | Buccal Bias | **1** | 1 | 1 | 1 |
| Lbi | Lingual Bias | **1** | 1 | 1 | 1 |
| Rad | Radius of Initial Conditions | **2** | 2 | 2 | 2 |
| Deg | Protein Degradation Rate | **0.075** | **0.1** | **0.1** | **0.1** |
| Dgr | Downward Growth | **10000** | 10000 | 10000 | 10000 |
| Ntr | Nucleus Traction | **0.00001** | 0.00001 | 0.00001 | 0.00001 |
| Bwi | Border Width | **0.7** | 0.7 | **0.9** | **0.9** |
| Ina | Initial Activator Concentration | **0** | 0 | 0 | 0 |

**Supplementary Table S2.** ToothMaker parameters set for catshark tricuspid tooth, with modifications performed to replicate increasing cusp number in development.
