## Supplemental Table 3 for "An ancient dental signalling centre supports homology of an enamel knot in sharks"

| *Symbol* | *Parameter* | *Seal* | *Shark* |
| --- | --- | --- | --- |
| Egr | Epithelial Growth | 0.022 | 0.0225 |
| Mgr | Mesenchymal Growth | 200 | 200 |
| Rep | Cell Repulsion | 1 | 1 |
| Swi | Distance from 0 where borders defined | 0 | 0 |
| Adh | Cell Adhesion | 0.001 | 0.001 |
| Act | Activator Self-regulation | 0.2 | 1 |
| Inh | Inhibitor Strength | 1.5 | 13 |
| Sec | Secondary Signals | 0.03 | 0.03 |
| Da | Activator Diffusion | 0.17 | 0.2 |
| Di | Inhibitor Diffusion | 0.2 | 0.2 |
| Ds | Growth factor diffusion Rate | 0.2 | 0.2 |
| Int | Initial Inhibitor Threshold | 0.15 | 0.19 |
| Set | Secondary Signal Threshold | 0.95 | 0.95 |
| Boy | Mesenchymal Buoyancy | 0.1 | 0.1 |
| Dff | Differentiation Rate | 0.00045 | 0.0004 |
| Bgr | Border Growth | 1 | 1.5 |
| Abi | Anterior Bias | 15 | 7.5 |
| Pbi | Posterior Bias | 14 | 7.5 |
| Bbi | Buccal Bias | 1 | 1 |
| Lbi | Lingual Bias | 1 | 1 |
| Rad | Radius of Initial Conditions | 2 | 2 |
| Deg | Protein Degradation Rate | 0.06 | 0.075 |
| Dgr | Downward Growth | 15000 | 10000 |
| Ntr | Nucleus Traction | 0.00001 | 0.00001 |
| Bwi | Border Width | 0.8 | 0.7 |
| Ina | Initial Activator Concentration | 5 | 0 |

**Supplementary Table S3.** ToothMaker parameters set for catshark tricuspid tooth, in comparison with seal-like mammalian tooth.
