## Supplemental Table S4 for "An ancient dental signalling centre supports homology of an enamel knot in sharks"

| Gene | Forward Primer | Reverse Primer |
|---|---|---|
| *lef1* | CATGCACTCTACAGGGATCCC | TCTGGATCAGAGTCTTGCTGC |
| *β-catenin* | AGTGGTTAAGCTACTGCACCC | AAGCTAGCATCATCTGGACGG |
| *shh* | TGACTCCCAATTACAACCCGG | TCAGGTCCTTCACTGACTTGC |
| *mdk* | GACAGGGTCCTCTGAAGCTG | TTAGGGTTCCATTGCGAGTC |
| *fgf3* | CTTGTTGCTGAGTCTTCTGGC | AACTCTTCAGCAGGTTCTCCC |
| *fgf10* | TGGATACTGACAAAGGGTGCC | GACATCGTGTCTCACCACTATTGG |
| *wnt11* | TCTGACATGAGGTGGAACTGC | TCTCTTGAGTTCCGTTGGAGC |
| *dkk1* | TGCCTCTACAATGTCGTGAGC | GTGCAGCCTCGAATTCTTGC |
| *bmp4* | GGAGCACAGGTCTATGGAAAGG | GGAGCACAGGTCTATGGAAAGG |
| *smad1* | GGAATCCGAGACACTCTTGGC | TTCAACAACCAGCTCTTCGCG |
| *smad3* | TAGTCACCATGAGCTTCGAGC | CCAATGTGCCTTCTTGTCAGC |
| *isl1* | ATTGTTCGGGACTAAATGCGC | TGCAGCGTTTGTTCTGAAACC |
| *jag1* | GGGCGACACTATAGAGAAGGC | ACAGGGATCAGAGATGCAAGC |
| *jag2* | AGCTGTACTGTGGCAATCTCC | GGATGCAACTGCTGTTCTTCG |
| *bambi* | GCATCTAACTGTGTGGCAACG | TCCAAGTCTAACTTCGCCACC |
| *sfrp3* | CCGTCATGAGGAGGTACAACC | TTCTGTTCCTCTGCTTCGACG |
| *smad7* | TCCTTGCCGGTACTGATATGC | GTGTGAAATCGTGGTCGTTGG |
| *runx2* | ATCTCTCAATCCTGCACCAGC | CCAGACAGACTCATCAATCCTCC |
| *wwtr1* | AGTTCCAGGTTCAGCACATCC | GCATTAGGTCCTCGCCTTCC |
| *foxl1* | TCAGAGGGTGACATTGAACGG | CTGATGGAGAAGGGTTGGACC |
| *foxl1l* | GTATCTCGACCTGCCTACAGC | ATGTCAGATGCCCAGTCTTGG |
| *notch2* | AGAATGGAGGCACTTGTCAGG | AGCCTCCTTTGTTCAGACAGG |
| *msx2* | TCACCGAAGTATTGTGCCTCC | GGAGCACAGGTCTATGGAAAGG |
| *axin2* | GACGGACAGTAGCGTAGATGG | TGGTGGATGTGATGATGGTGG |
