## Supplementary figures and images for "An ancient dental signalling centre supports homology of an enamel knot in sharks"

### Supplemental Figure S1

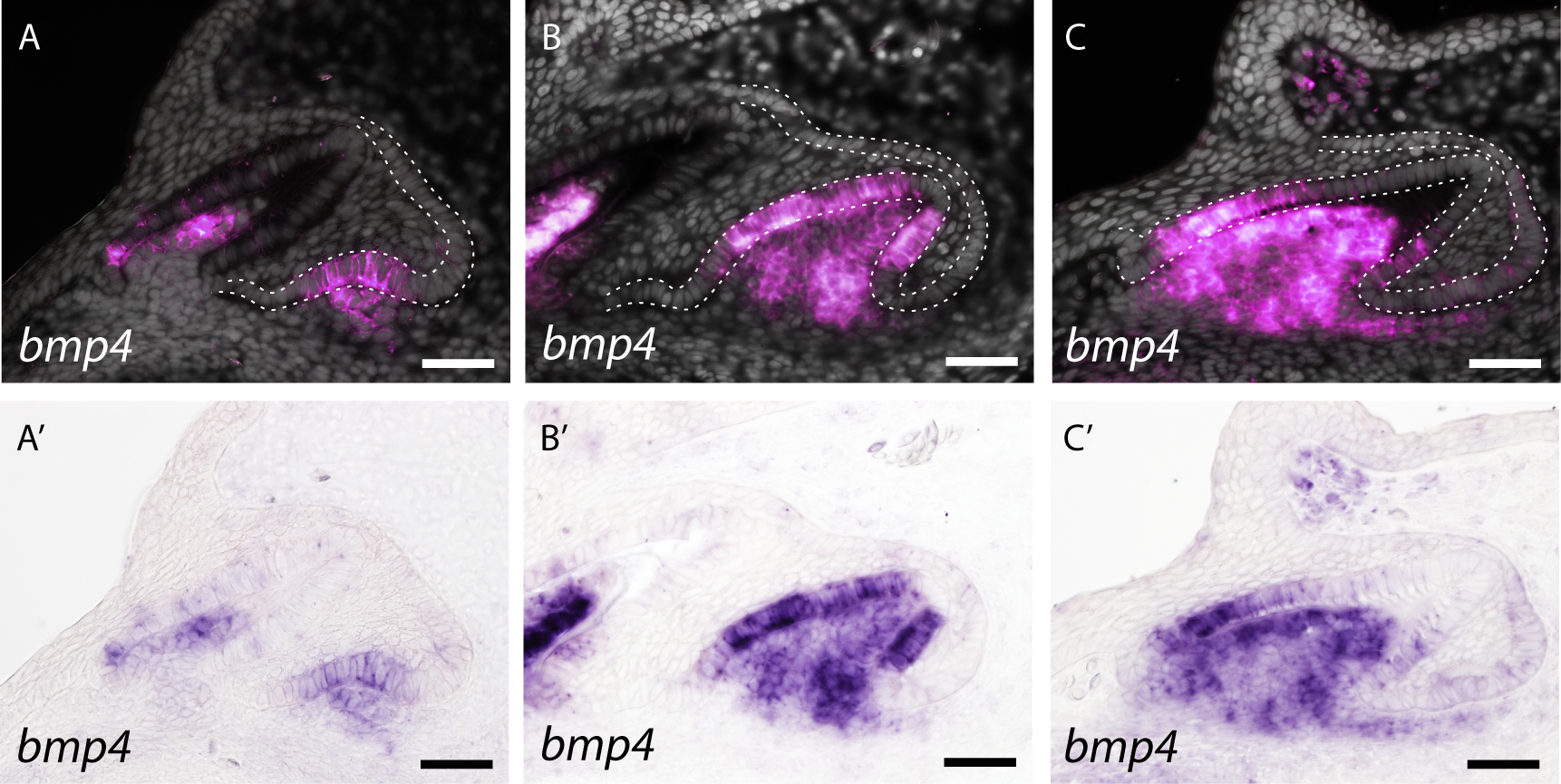

### Supplemental Figure S2

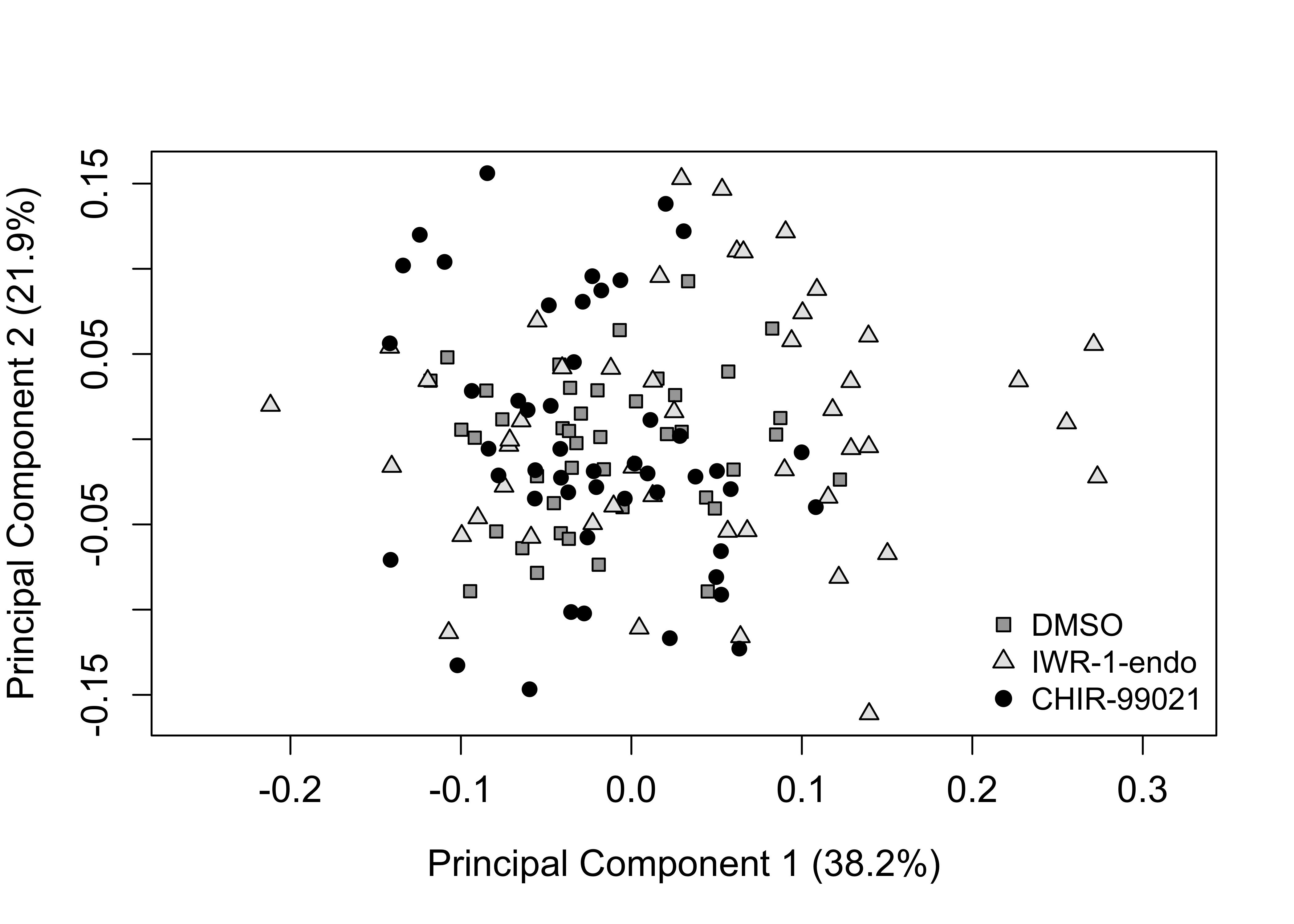

### Supplemental Figure S3a

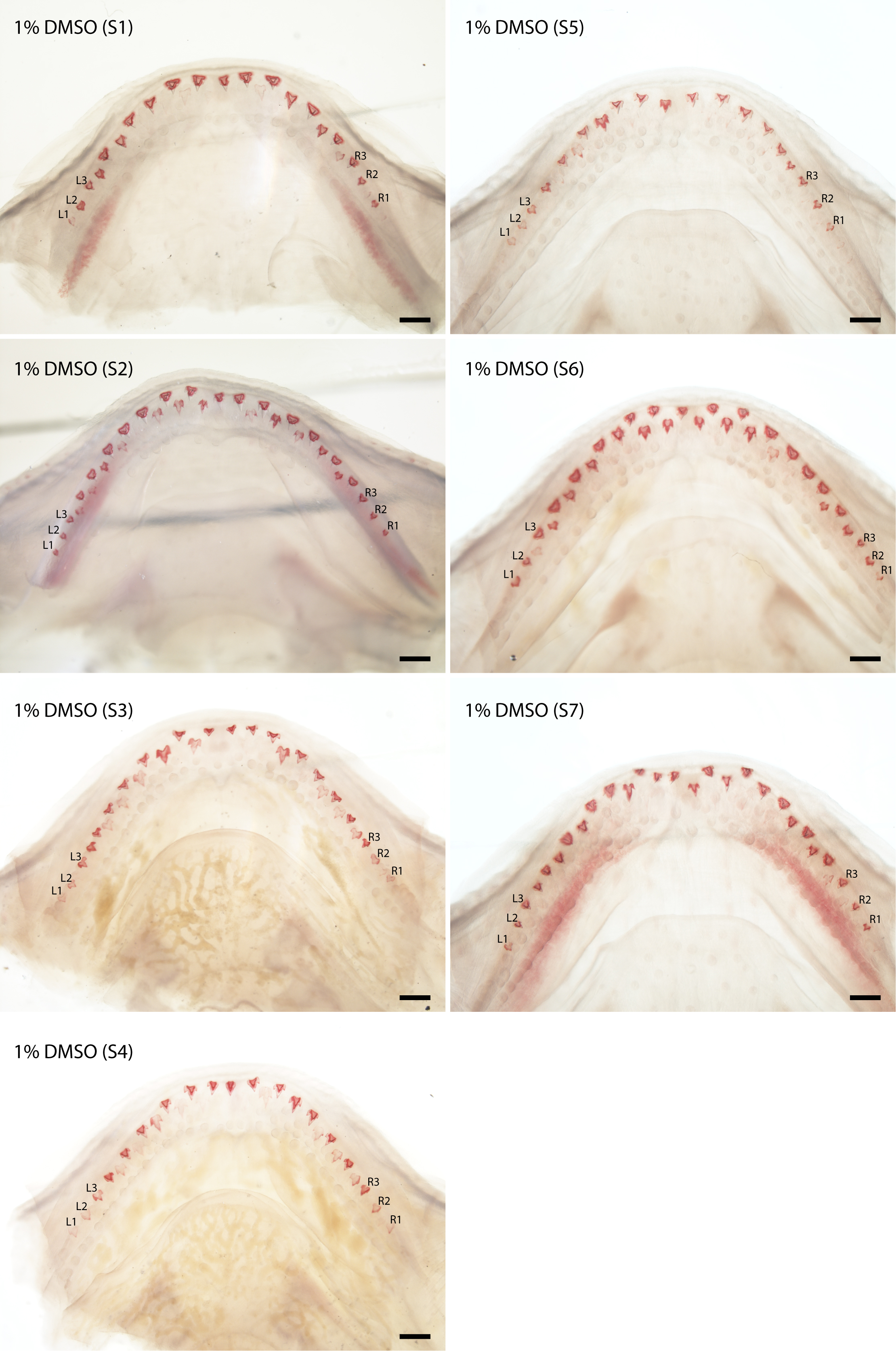

### Supplemental Figure S3b

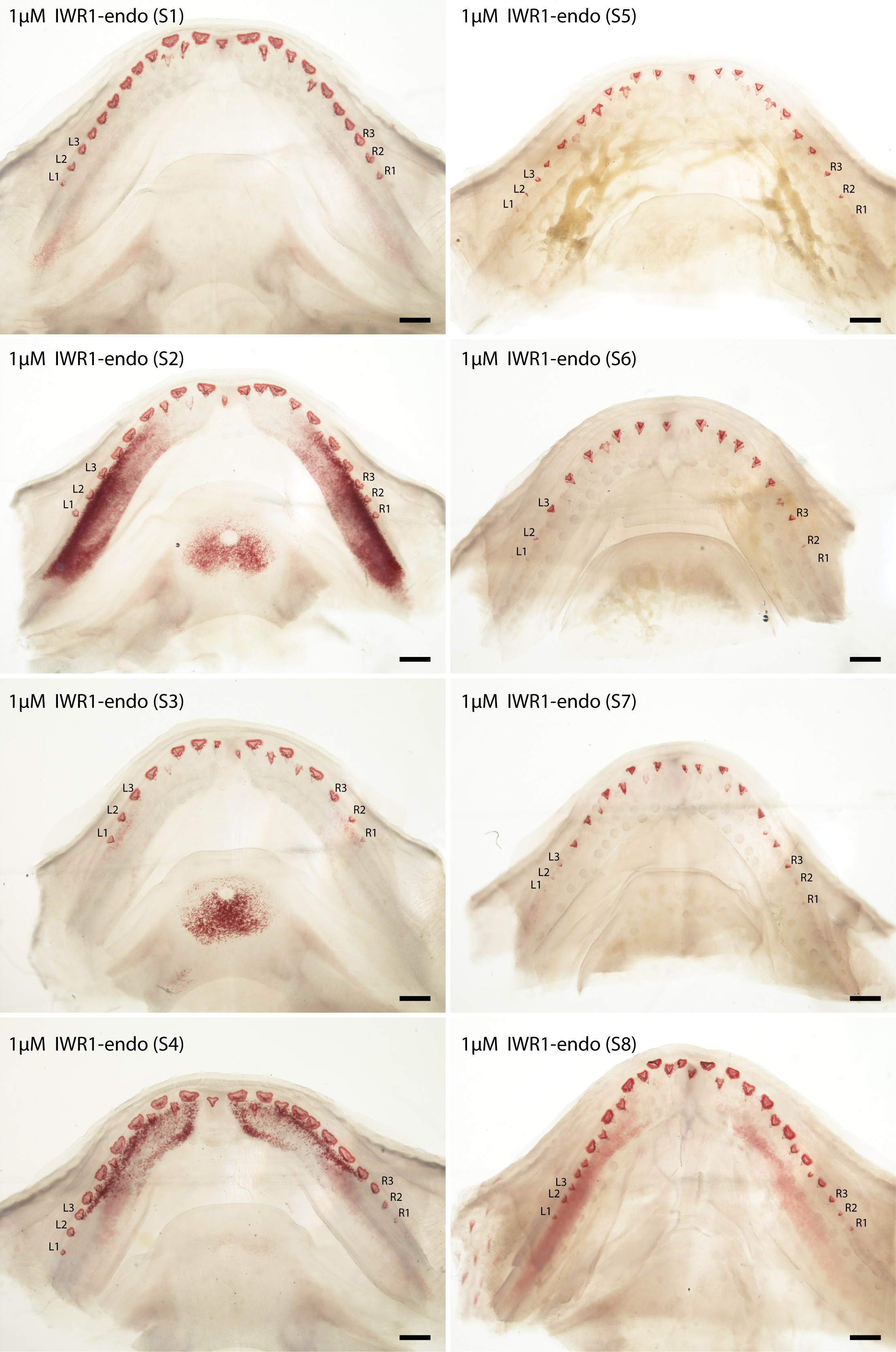

### Supplemental Figure S3c

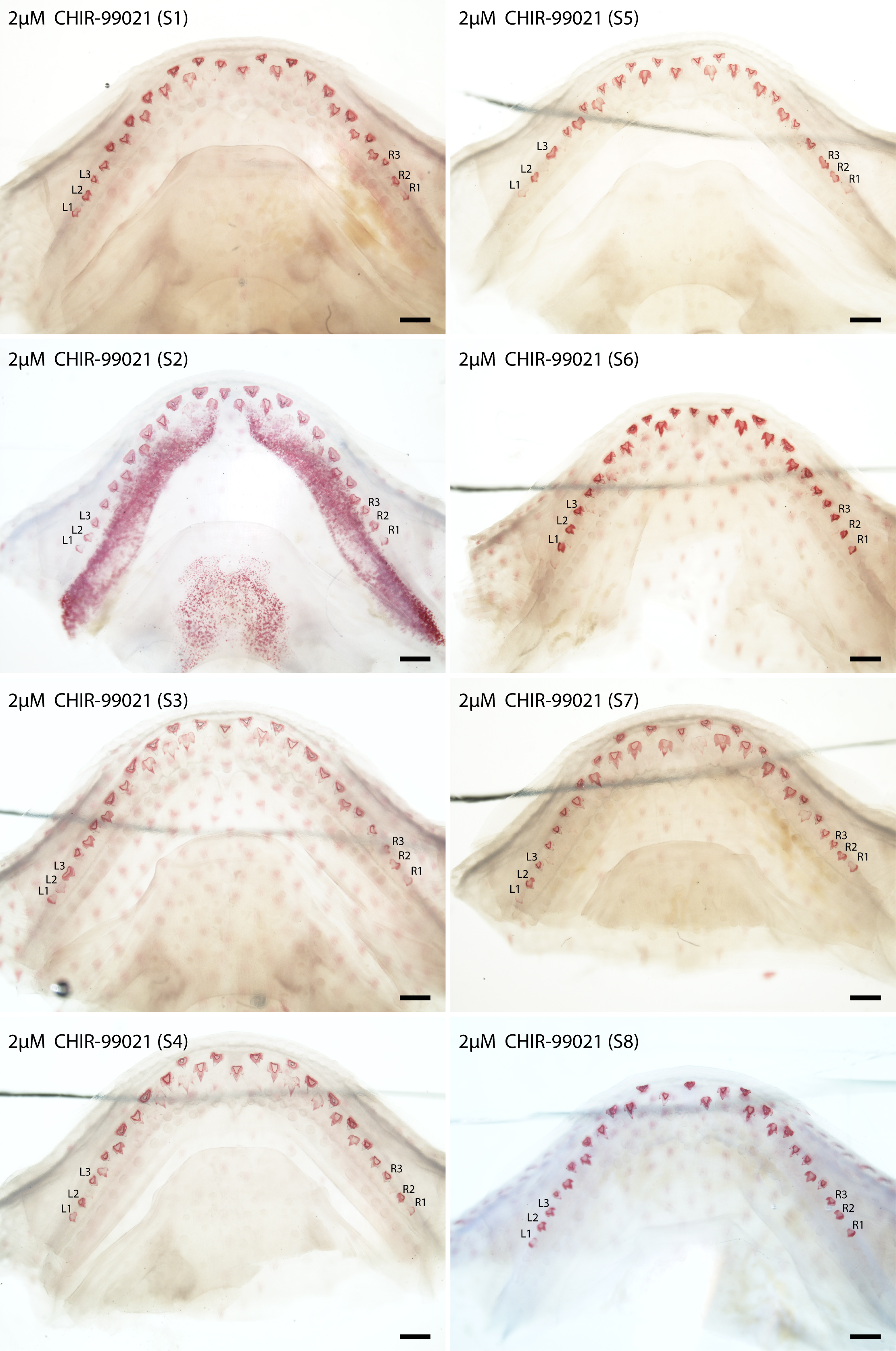

### Supplemental Figure S4

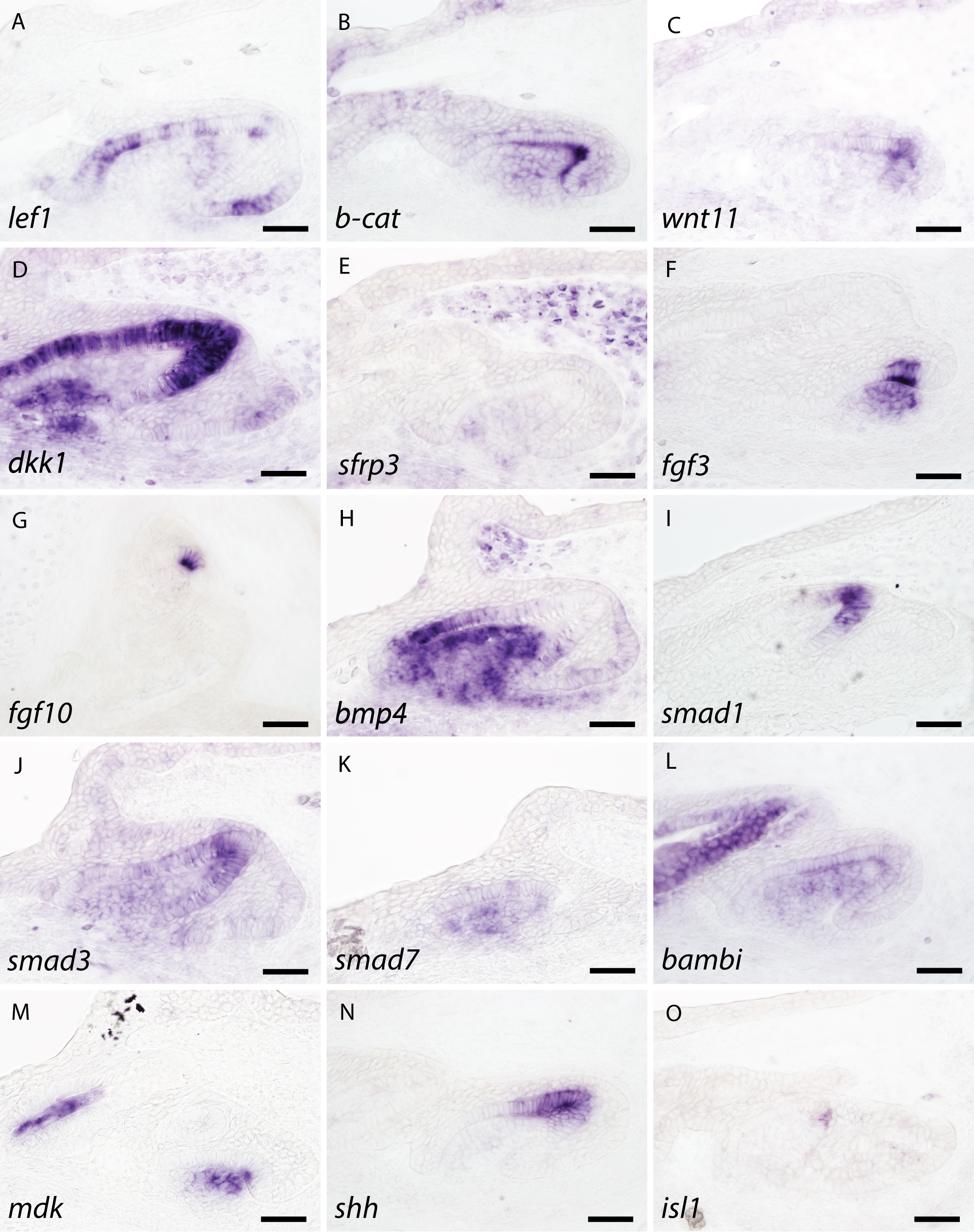
